# High-Fidelity Long-term Whole-embryo Lineage and Fate Reconstruction by Iterative Tracking with Error Correction

**DOI:** 10.64898/2026.03.12.711203

**Authors:** Mengfan Wang, Qinghua Zhang, Congchao Wang, Yunfeng Chi, Wei Zheng, Zeyu Mu, Xiangyu Cao, Weizhan Zhang, Boao Yang, Alexander F. Schier, Joaquin Navajas Acedo, Yinan Wan, Guoqiang Yu

**Affiliations:** Department of Automation, Tsinghua University; Beijing, 100084, China; Bradley Department of Electrical and Computer Engineering, Virginia Polytechnic Institute and State University; Arlington, VA 22203, USA; IDG/McGovern Institute for Brain Research, Tsinghua University; Beijing, 100084, China; Beijing National Research Center for Information Science and Technology; Beijing 100084, China; Biozentrum, University of Basel; Basel, 4056, Switzerland; Allen Discovery Center for Cell Lineage Tracing, Seattle; Washington, 98195, United States

**Keywords:** lineage reconstruction, fate mapping, cell tracking, embryonic development analysis

## Abstract

Reconstructing the complete cell lineages and fate maps of a living embryo has remained a central challenge in developmental biology. Here we introduce ITEC (Iterative Tracking with Error Correction), a fully unsupervised method that automatically reconstructs the lineage of every cell in the embryo with high fidelity. ITEC was validated with manually annotated lineages on four cross-species datasets including zebrafish, mouse, and *Drosophila*. We reconstructed the developmental lineages of a zebrafish embryo from terabyte-scale data of total 18.5 million cells with an estimated accuracy of over 99.7%. With ITEC, we revealed spatiotemporal dynamics of key morphogenetic processes such as somite boundary formation, demonstrated the various patterns of spatial sorting among adjacent organs or anatomical regions, and identified associations between cellular movement and spatial transcriptomics. ITEC provides a powerful platform for retrospective fate mapping and systematic exploration of developmental dynamics at the cellular scale and single-embryo level.

## Introduction

Reconstructing the complete developmental lineage for every cell as a living embryo develops has long been considered a “holy grail” of developmental biology (*1–4*). Achieving this goal is critical for understanding how cells interact to form complex tissues and organs, how spatial patterning emerges during development, and how cellular diversity arises from initially homogeneous populations (*5–9*). With the advancement of microscopic imaging technology (*3, 10–17*), long-term and large-scale multidimensional datasets continue to emerge to record the dynamics of the whole embryo at the single cell level (*18*), establishing the foundation to achieve this goal. However, manual analysis of the whole embryo is infeasible for such terabyte-scale datasets with tens of millions of cells. Conversely, a selective manual annotation of a few cells is prone to bias and loses the systematic view of the whole embryo. Therefore, solving the bottle-neck of automatic and comprehensive lineage reconstruction would be a major advance in developmental biology.

Since the emergence of whole-embryo dynamic imaging data, there have been several efforts to computationally reconstruct the lineages and map cell fates, through either adopting generic cell tracking techniques (*19–37*) or designing new approaches for embryonic data (*1, 38–41*). Among these are three state-of-the-art methods, TGMM (*1, 41*), Linajea (*41, 42*), and Ultrack (*38, 43*). However, major challenges remain mainly due to two requirements: First, an extremely high accuracy is required for both detecting cells at each time point and linking cells at two adjacent time points, because any error in a reconstructed lineage along hundreds of time points will result in an erroneous, useless or misleading reconstruction. Second, the embryonic data is heterogeneous with low signal-to-noise ratio, which is a consequence of the variations in the transmissive properties of embryos, and the experimental necessity of biologically relevant conditions for embryo development.

Here, we present a full-process embryonic tracking pipeline, ITEC (Iterative Tracking with Error Correction), which achieves lineage reconstruction of whole embryos with an unprecedented accuracy (Fig. 1 and Fig. 2). ITEC exploits the special properties of the embryonic imaging data and builds upon our recent development of a new machine learning technique for data association (*44*). Though the heterogeneity and low signal-to-noise ratio of embryonic imaging data inherently lead to imperfections in initial cell detection and linkage, long-term spatiotemporal information precisely provides strong support to correct errors and pursue ultra-high accuracy. We developed a strategy of iterative error correction, which uses the tracking results as feedback to correct cell detection errors, and updates the tracking results based on the corrected cell detection, ultimately forming a stable closed-loop system (Fig. 3). Errors decrease continuously with iterations, which significantly improves the accuracy and robustness of tracking, particularly in scenarios with high imaging heterogeneity and low image clarity. The concept of iteratively correcting errors seems intuitive, but it was not clear how to identify and correct errors. It is our newly invented and record-setting minimum-cost circulation model (*44*), that enables the implementation of error correction. ITEC is user-friendly as well, as it adopts fully unsupervised techniques and requires no data labelling or human annotations. High-precision cell detection and tracking can be achieved by simply adjusting a small number of biologically meaningful parameters.

**Fig. 1.**
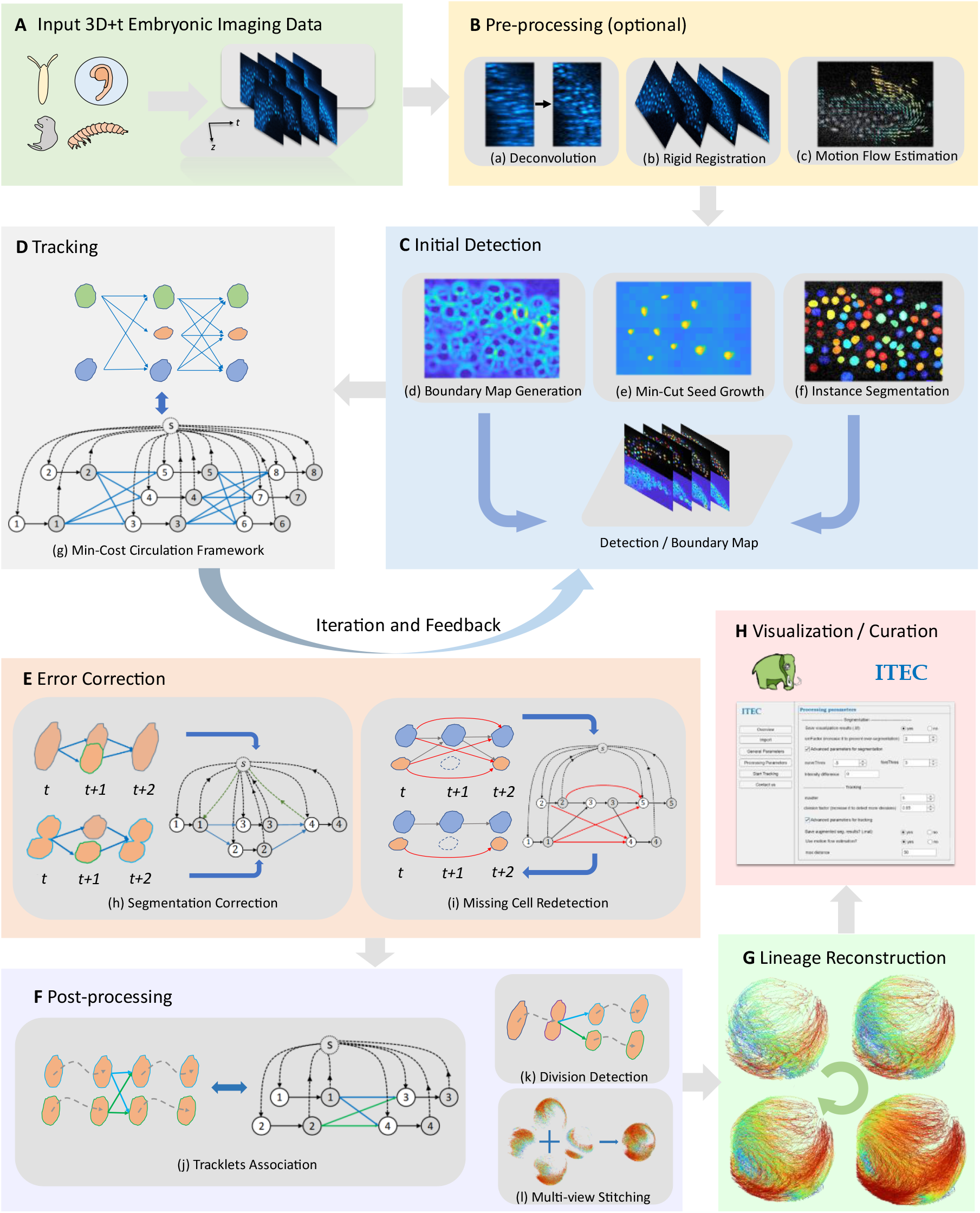
Flowchart and pipeline of ITEC. (A) A typical 3D+t input data, where each time frame consists of 3D image stacks acquired from light microscopy. ITEC supports embryonic development data of multiple species. (B) Pre-processing, which includes optional deconvolution, rigid registration, and motion flow estimation. Users can also perform individual processing steps tailored to their specific data. (C) Initial detection, utilizing high-order image information (such as in (d)) and the min-cut algorithm (such as in (e)) to achieve pixel-level instance segmentation (such as in (f)) of cells. (D) Tracking, where the problem of cell linkage across frames is modeled as a maximum a posteriori estimation problem and solved using a minimum-cost circulation framework. (E) Error correction, which includes segmentation correction module and missing cell redetection module, used to correct errors through feedback based on the last round of tracking results. (F) Post-processing, which includes tracklets association, division detection, and multi-view stitching, further enhancing the accuracy and completeness of cell linkage. (G) Lineage reconstruction. Lineages are extracted from tracking results and reformatted for the downstream analysis. (H) Visualization and curation. ITEC supports parameter tuning and execution through visualization tools, and allows visualization and curation of tracking results using Mastodon (*47*).

**Fig. 2.**
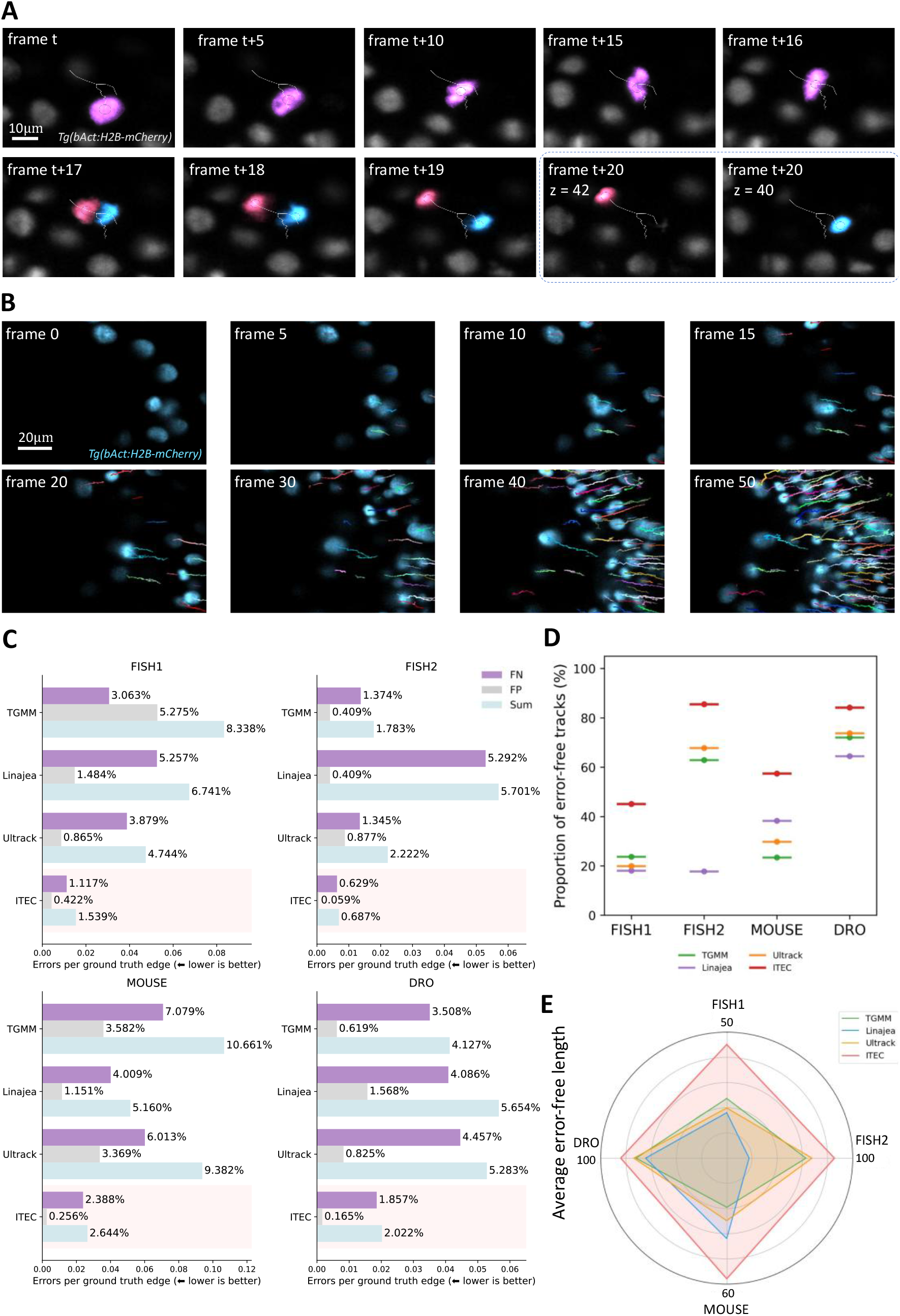
Tracking and lineage reconstruction performance of ITEC on four cross-species datasets. (A) Illustration of a typical cell track from ITEC. It consists of 20 frames. The pink cell divided in frame 17. (B) Tracks of cells from ITEC under crowded conditions. Tracks of different cells are represented in different colors. (C) Errors per ground truth edge of ITEC and peer methods. ITEC achieves the best performance with a large margin across all four datasets. (D) Proportion of error-free tracks of ITEC and peer methods. This metric represents the proportion of completely correct lineages to the total annotated lineages. (E) Average error-free length of ITEC and peer methods. This metric is the number of consecutive time points over which there are no tracking errors.

**Fig. 3.**
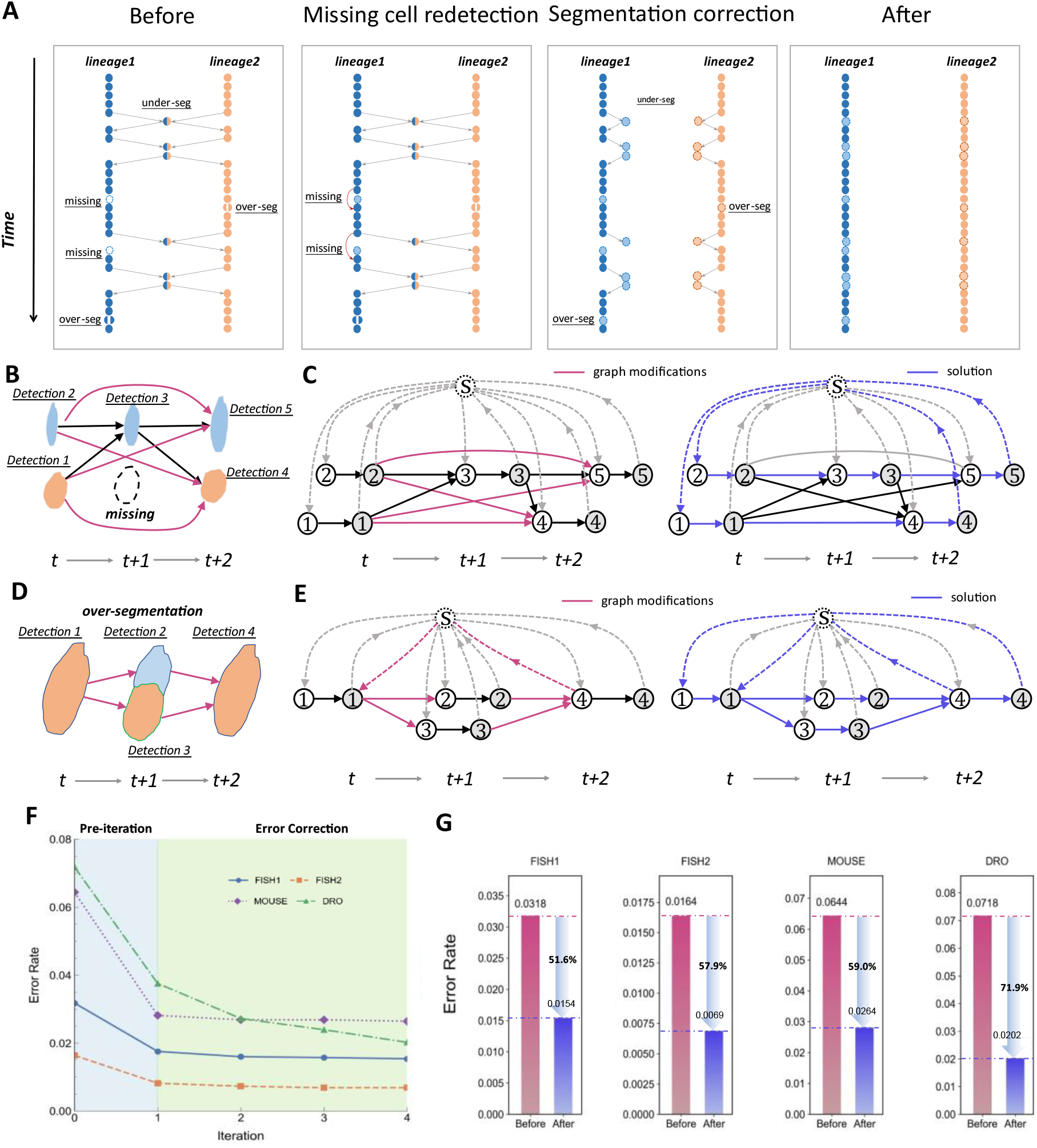
ITEC fully utilizes spatiotemporal information to iteratively correct errors. (A) Illustration of segmentation improvement through error correction module. Initially, there are 10 errors in these two lineages, including 2 missing, 2 over-segmentation, and 6 under-segmentation errors. After missing cell redetection, 2 errors are fixed, and the remaining 8 errors are resolved through segmentation correction, resulting in continuous and correct tracks. (B) Illustration of missing cell error. There are two tracks in (B). The first one is composed of detection 2, 3 and 5 and the second one consists of detection 1 and 4. A detection is missed in the second track at frame t + 1. (C) The network that allows cells to jump over some frames to be compatible with missing cells due to errors cell detection. This network design legitimates the tracking to continue from frame t to t + 2 by allowing linkages between detections beyond adjacent frames. (D) Illustration of over-segmentation error. In frame t + 1, a cell is wrongly split into two detections, while the segmentation is correct in frame t and t + 2. (E) The new design of the network allows detections to split and merge. For each detection in the (D), it is represented by two nodes in the network shown in (E). Both detection 2 and 3 can link to detection 1 or 4. With such design, the over-segmentation cells can be processed. (F) Curve of error rate vs. number of iterations. Before iteration, the error rates of FISH1, FISH2, MOUSE and DRO datasets are 0.0318, 0.0164, 0.0644, and 0.0718, respectively. After the first round of iteration, the error rates become 0.0175, 0.0082, 0.0281, and 0.0376, respectively. (G) The error rate decreased after completion of iteration. In the FISH1, FISH2, MOUSE and DRO datasets, the iterative strategy reduced the error rates by 51.6%, 57.9%, 59.0% and 71.9%, respectively.

With the aid of ITEC, we automatically reconstructed lineages of over 18.5 million cells cumulatively across the entire zebrafish embryo at terabyte scale (Fig. 4). With an evaluation of a randomly selected subset of lineages spanning up to 600 time points and including over 30,000 cells, ITEC achieved an unprecedented high accuracy of over 99.7% per cell linkage and improved the lineage accuracy to 48.5% from 20% of the best peer methods. On four benchmark datasets, including two zebrafish, one mouse, and one *Drosophila* embryo development time-lapse datasets, ITEC outperformed all the three state-of-the-art methods mentioned above by a large margin across all metrics (Fig. 2). Particularly, ITEC reduced the errors per ground-truth cell linkage by at least 51.6% compared to the second-ranked method on any benchmark dataset.

**Fig. 4.**
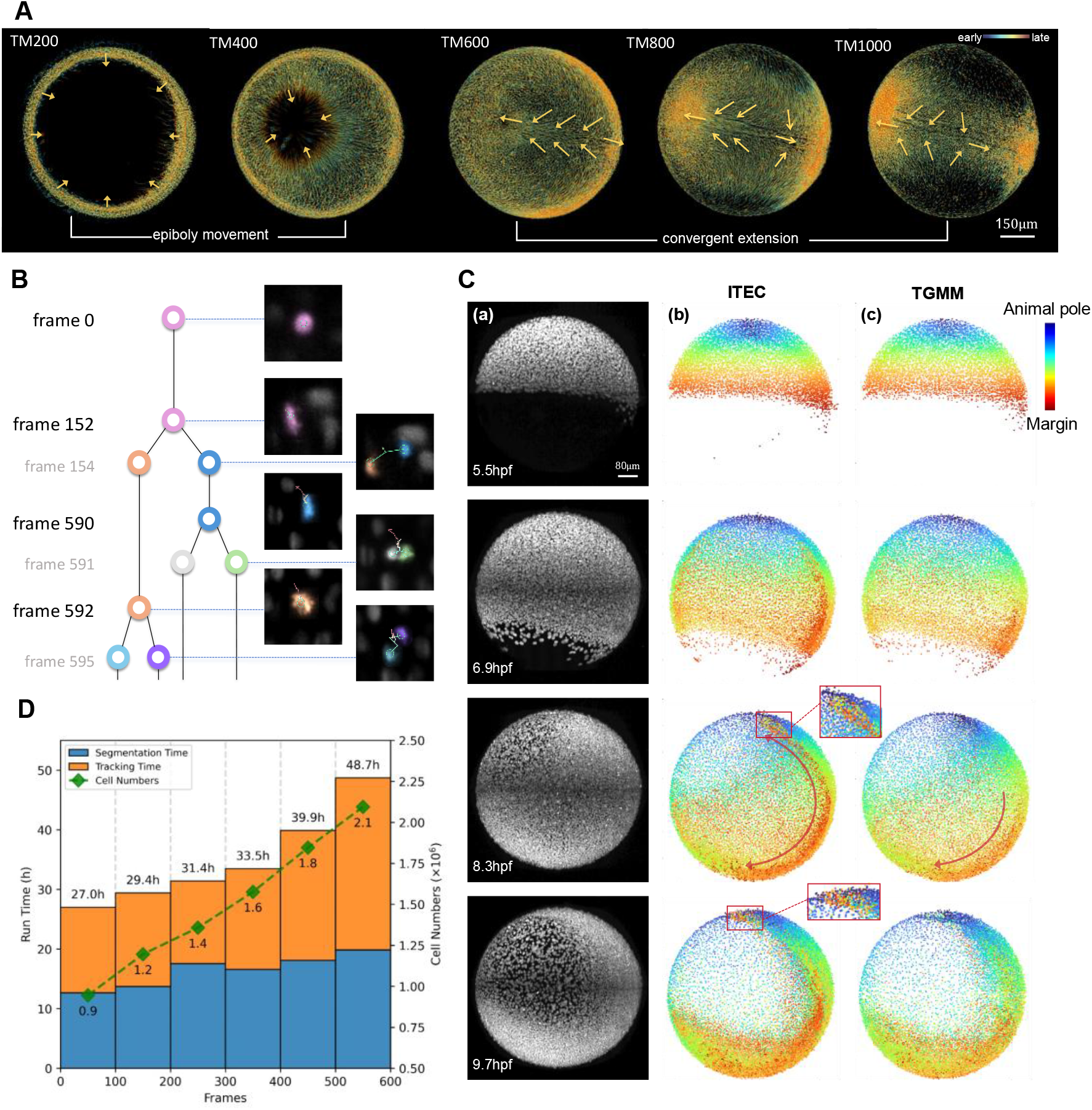
ITEC enables long-term lineage reconstruction at terabyte level. (A) Screenshots of long-term zebrafish lineage reconstruction from the Mastodon visualization platform. In the first half, cells primarily undergo epiboly movement, while in the latter half, cells undergo convergent extension. The arrows indicate the cell motion directions. (B) A typical lineage tree from ITEC. The ancestor first splits once at frame 152, and the two offspring cells split twice at frames 590 and 592, respectively. Our method can capture three divisions and accurately track the migration of all children. (C) Snapshots and the reconstruction comparison of the dataset FISH2. (a) The max projection of datasets. The rows from the first to the last show the progress of the zebrafish embryo at the time point 0, 240, 480, and 720, corresponding to 5.5, 6.9, 8.3, and 9.7 hpf, respectively. (b) The reconstruction results of ITEC. Every dot represents one detected cell in the corresponding time point, and the dot color represents the initial cell position at the time point 0. A blue dot indicates a cell originates near the animal pole, while a red dot tells the cell originates at the near the marginal zone. The purple arrows at 8.3 hpf indicates the hypoblast and the red box at 9.7 hpf indicates the hatching gland. (c) The reconstruction results of TGMM according to the same arrangement as (b). (D) Runtime variation in relation to the count of cells and frames. ITEC has an option of parallel computing to enable the distributed processing of TB-level data.

We applied ITEC to analyze the dynamic features of embryogenesis. During somite formation in zebrafish, we found that sharp somite boundaries gradually and irreversibly emerge from an initially intermixed population of progenitor cells (Fig. 5). During gastrulation, we characterized the diversity of cell mixing and rearrangements during processes such as convergence and midline formation (Fig. 6). To decipher the molecular mechanisms underlying cell behaviors, we combined embryonic lineage reconstruction with our recently developed whole-mount spatial transcriptomics (*9*) (Fig. 7). In summary, ITEC transforms cell tracking from an error-prone, fragmented process into a robust end-to-end solution, thereby turning large-scale and long-term embryo lineage reconstruction from a burdensome task into a practical and routinely feasible approach for biological research.

**Fig. 5.**
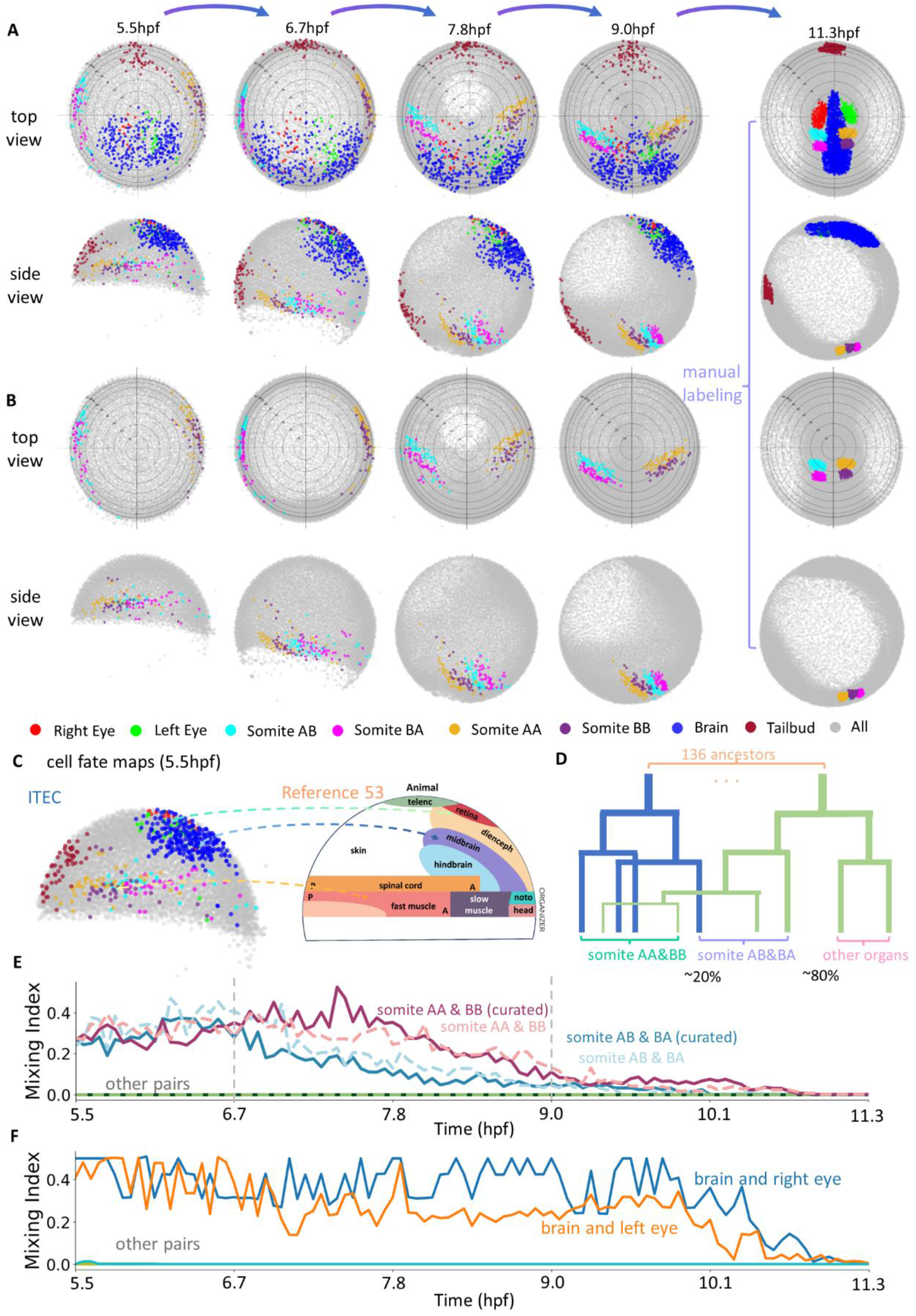
Fate mapping and mixing index analysis of zebrafish embryo development dataset based on ITEC. We manually segmented four classes of zebrafish organs: eyes, brain, somites, and tail bud, including the left and right eyes and four somites AA, AB, BA and BB, at the early organogenesis stage (11.3 hpf, the last time point of the data). (A) The top view and side view of the fate mapping results except for the last column, which visualizes the manual segmentations. The first column to the last column shows the fates of the organs at the time point 0, 200, 400, 600, and 1000, corresponding to 5.5, 6.7, 7.8, 9.0, and 11.3 hpf, respectively. (B) The top view and side view of the somite fate map according to the same arrangement as (A). (C) Comparison of a fate map of the zebrafish embryo at the start of gastrulation from (*53*) with the fate map generated by ITEC. (D) Schematic of ancestral and descendant lineage tree for somites AA, AB, BA and BB. (E) Mixing index for the four somites with and without curation. The ancestors of the somite AA&BB and somite AB&BA pairs gradually separate over time, whereas all other pairs such as AA vs AB, and BB vs BA remain completely unmixed. (F) Mixing index for various organ pairs in the zebrafish embryo. The ancestors of the brain/left eye and brain/right eye pairs gradually separate over time, while other pairs show almost no mixing throughout development.

**Fig. 6.**
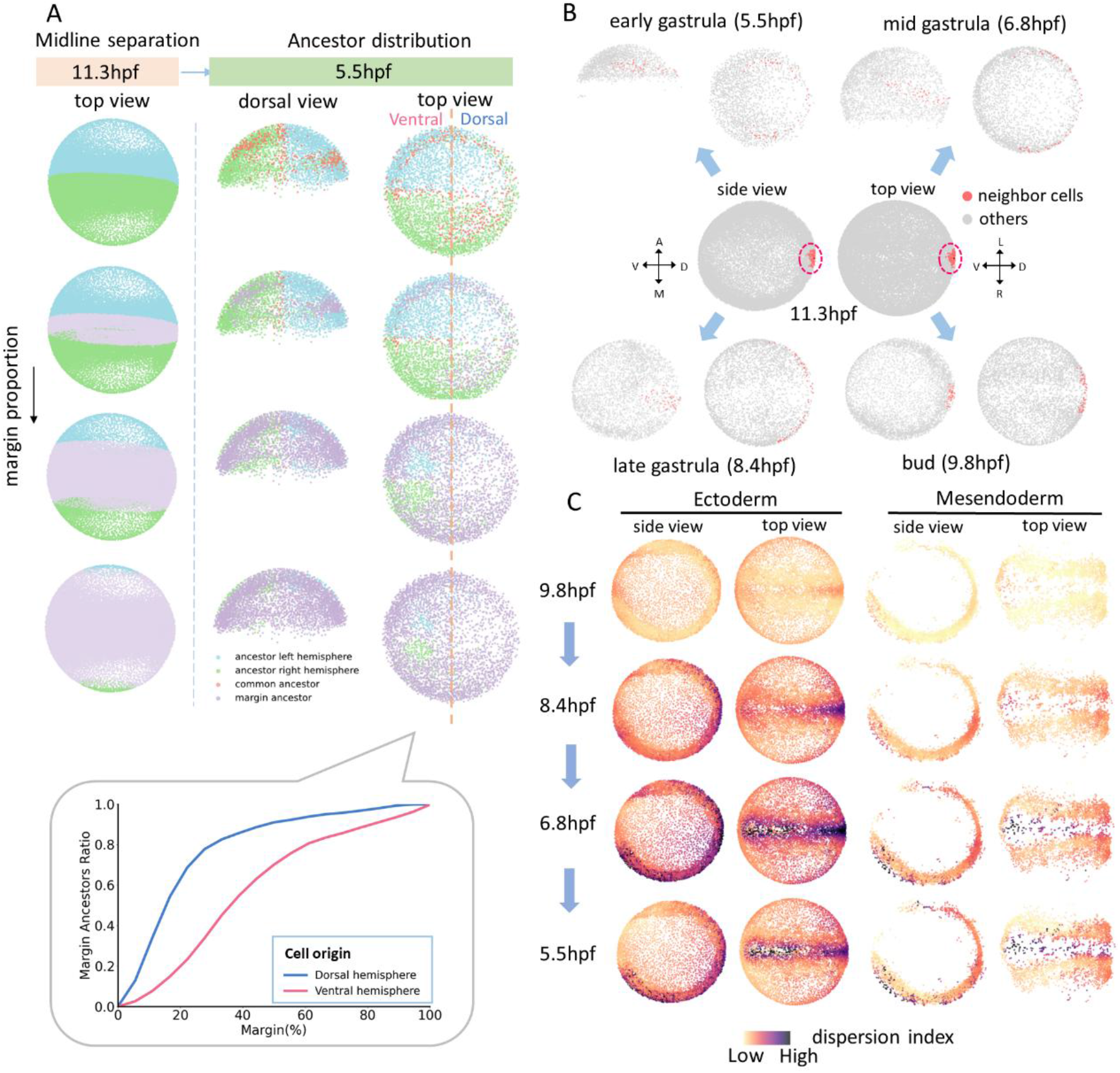
Analysis of midline formation and cell dispersion during zebrafish embryonic development. (A) Cell lineage tracing of cells bilateral to the midline in zebrafish. It shows the origin of cells bilateral to the midline and in the margin as the margin proportion increases. The bottom subfigure shows the proportion of ancestors in the margin region relative to all ancestors on the dorsal and ventral sides, respectively. (B) Ancestral distribution of a cell population at 11.3 hpf traced back to 5.5, 6.8, 8.4, and 9.8 hpf, respectively. Ancestors are represented by red scatter points. (C) Mean distance dispersion index of cells at 11.3 hpf traced back to their ancestors at 5.5, 6.8, 8.4, and 9.8 hpf, respectively. A darker color in the plot indicates a more discrete distribution of the ancestral origin of cells in that area.

**Fig. 7.**
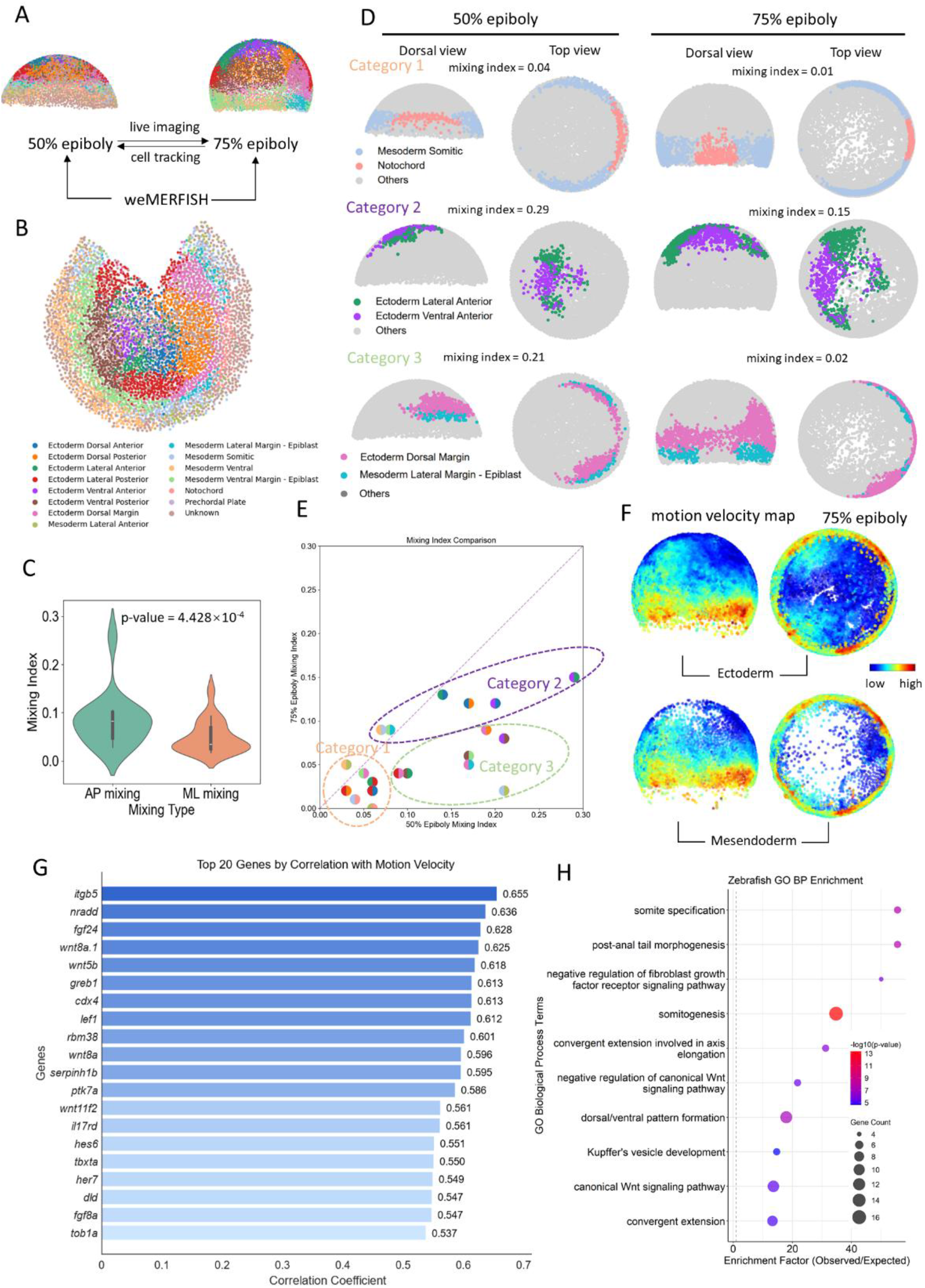
Integration of cell tracking with spatial transcriptomics weMERFISH. (A) Schematic diagram of the integration of live imaging and spatial transcriptomics. Cell association from 50% epiboly to 75% epiboly is achieved through cell tracking via live imaging, while the matching between live imaging and spatial transcriptomics is accomplished using the global optimal mapping method. (B) Distribution of the 14 regions defined by gene expression profile at 75% epiboly. (C) Wilcoxon test for cell mixing index along the anterior-posterior (AP) and medial-lateral (ML) axes. (D) Comparison of the mixing index of different mixing patterns between 50% epiboly and 75% epiboly. (E) Distribution of the mixing index at 50% epiboly and 75% epiboly. Based on the scatter of the points, we clustered them into three categories. (F) Cell migration velocity of ectoderm and mesoderm at 75% epiboly. (G) Top 20 genes most strongly and positively correlated with migration speed and their correlation coefficients. (H) GO term enrichment analysis of top 90 genes positively correlated with migration speed.

## Results

### Unsupervised full-process pipeline of Iterative Tracking with Error Correction (ITEC)

Long-term tracking of embryonic cells demands extreme precision, imposing high requirements on the collaboration between processes such as cell segmentation and cell linkage. Iterative Tracking with Error Correction (ITEC) is a high-fidelity and full-process cell tracking pipeline designed for embryonic lineage reconstruction and fate mapping, which is potentially applicable for the generic cell tracking tasks but not comprehensively tested yet. It is an end-to-end process, covering the entire procedure from raw data pre-processing to the final reconstruction of lineages, with no user intervention required in between. In addition, it is entirely based on unsupervised learning and does not require any labeling. For more technical details, please refer to Online methods.

The flowchart of ITEC is shown in Fig. 1. First, we provide users with optional pre-processing operations to improve image quality and correct imaging artifacts, including deconvolution (*45*), rigid registration and motion flow estimation for non-rigid deformation (*27*) (Fig. 1B). These steps allow meaningful biological signals to be enhanced from imaging data with heterogeneous cell activity limited by noise and resolution. Then we perform initial cell detection by applying principal curvatures to mimic human perception (Fig. 1C, Fig. S2 and Fig. S4) (*46*). We combine curvature maps under different scale filtering and utilize robust statistical tests to effectively exclude background regions and counteract data heterogeneity. The boundary refinement problem is modeled as the min-cut graph problem to ensure pixel-level instance segmentation. After the initial detection, we transform the cell linkage problem into a global discrete optimization problem and build the minimum-cost circulation framework that can efficiently handle millions of cells (Fig. 1D).

Subsequently, we use an iterative error correction strategy to further refine the segmentation and cell linkage results based on the existing tracking results, making full use of the rich spatiotemporal information in long-term lineage reconstruction (Fig. 1E). This strategy involves multiple iterations, and results gradually become stable in the process of segmentation correction and recovering the missing cells. To ensure the continuity of the tracks and consistency with biological hypotheses, we perform tracklets association, division detection, and multi-view stitching (Fig. 1F). For user convenience, we also design an interactive interface and integrate our tracking results with Mastodon (*47*), a Fiji (*48*) plugin for cell tracking analysis (Fig. 1H). Among all the technical innovations in developing the ITEC pipeline, the iterative error correction strategy and the minimum-cost circulation modeling of data association were the two most effective, which are further discussed and evaluated with ablation experiments in the next sections.

### ITEC outperformed peer methods on four cross-species datasets

To comprehensively compare our method with peer methods, we conducted experiments against ITEC and three state-of-the-art peer methods TGMM (*1, 39*), Linajea (*41, 42*), and Ultrack (*38, 43*). These methods respectively adopt statistical approaches, utilize deep learning techniques, and integrate segmentation and linking, reflecting the latest advancements in the field of cell tracking. More details about the peer methods can be found in Table S1. We then selected four cross-species datasets, including zebrafish, mouse and *Drosophila* embryo (FISH1, FISH2, MOUSE, and DRO) (Table S2) and constructed the ground truth by combining public annotations with our manual annotations. The annotations resulted in total of 44,555 ground-truth edge linkages. Characterized by diverse signal-to-noise ratio, temporal resolutions, spatial resolutions, and cell densities, performances on these datasets shall be a good indication of the robustness and generality of these tracking methods in real applications (Fig. 2A, Fig. 2B, Fig. S2A and Fig. S2B). For the definitions of the evaluation metrics, please see Fig. S1 and Supplementary Materials.

We first evaluated the accuracy of each edge linkage between adjacent frames. ITEC achieved a lead by a large margin across all four datasets (Fig. 2C). Its sum errors per ground truth edge (error rate) on FISH1, FISH2, MOUSE, and DRO were 32.44%, 38.53%, 51.24%, and 48.99% of that of the second-ranked method, respectively. Among these, ITEC gained a more significant lead on the two zebrafish embryo datasets FISH1 and FISH2, primarily due to the higher spatial continuity and stronger inter-frame correlation information. For the mouse embryo dataset, the opacity of mouse embryos and the long imaging intervals required for long-term imaging resulted in an error rate of ITEC exceeding 2.5%. Excluding ITEC, none of the three peer methods dominated the other two. TGMM worked well on FISH2 and DRO datasets, Ultrack suited for FISH1 dataset, and Linajea worked best in MOUSE dataset. Notably, during the evaluation process, we only need to set fundamental parameters such as grayscale, size, and filtering parameters, without making any modifications to the method ITEC itself. This indicates that our method exhibits high robustness and can meet the analytical needs of various model organisms used by the community.

While some lineages are difficult to reconstruct, leading to a lot of linkage errors, others are easier to follow, with small or no errors, and result in better lineage accuracy than expected through independence assumption. In order to quantify the performance on long-term lineage reconstruction, we selected *proportion of error-free tracks* and *average error-free length* to evaluate the continuity and reliability of tracks. Error-free tracks are complete paths along which cells can be followed correctly, while error-free length indicates the maximum number of frames a method can track correctly in a single, complete path. The reason for choosing these two metrics lies in the fact that biologists are interested in the tracking of cells during long-term migration. Thus, these metrics better reflect the feasibility of cell tracking that holds biological value compared to metrics such as edge linkage accuracy. We calculated the average of error-free length across all lineages in the ground truth to assess the continuity of the reconstructed lineages. In addition, we defined the proportion of error-free tracks out of the total number of lineages in the ground truth to measure the ability for long-term cell migration tracking. Experiments demonstrated that our method also achieved the best fidelity in lineage-scale evaluation (Fig. 2D, Fig. 2E and Fig. S3). In the FISH2 dataset, 85.5% of lineages remained completely accurate over 100 frames, compared to the next best of Ultrack being 67.7%. In the FISH1 dataset with 192 frames, longer imaging intervals and faster developing speed, ITEC achieved complete accuracy in 45.1% of lineages versus 23.9% obtained by the next best approach, TGMM.

### Iterative error correction strategy improves the overall tracking accuracy

A major challenge in high-fidelity lineage reconstruction lies in the accumulation of errors. On the one hand, from the perspective of long-term cell tracking, errors accumulate over time because one error in a long track can render the entire track invalid. On the other hand, errors can propagate within a certain spatial range, leading to a chain reaction of incorrect linkages between surrounding cells. The vast majority of these errors stemmed from the instability of segmentation and seemed to be unavoidable owing to the low and heterogeneous image quality. It is hard to achieve high image quality due to factors such as uneven illumination, variations in the transmissive properties of embryos, and limitations of imaging equipment (Fig. S2A). Moreover, to minimize toxicity, the experiments are often planned to have a signal-to-noise ratio as low as possible.

Interestingly, when annotating a time-lapsed image dataset, human experts can overcome these limitations by leveraging the rich spatiotemporal information residing in the data. Therefore, we designed an iterative error correction strategy, expecting an initial lineage reconstruction can be leveraged to identify and correct errors. For example, if there are cells at the same location in both frame *t* and frame *t*+2, there is a high probability of a cell being missed in frame *t*+1 (Fig. 3B). If there is only one cell in consecutive frames at a certain position, and in one frame it is detected as two cells, then that frame is likely to have undergone over-segmentation (Fig. 3D). We can leverage the cell segmentation and linkage results within a time window, combined with prior information such as cell size and spatial relationships, to infer the most probable scenarios, thereby compensating for missing cells or resolving segmentation errors. ITEC is capable of multiple iterations, in which the missing cell recovery and segmentation correction are performed sequentially in each iteration, gradually reducing errors during the iteration (Fig. 3A).

To validate the effectiveness of the iterative error correction strategy, we conducted ablation experiments: with only one round of error correction, the error rates of the FISH1, FISH2, MOUSE and DRO datasets were reduced by 45.0%, 50.0%, 56.4%, and 47.7% of the original error rates, respectively (Fig. 3F). After 3 iterations, the error rates tended to stabilize, and the final error rates on the four datasets were 1.54%, 0.69%, 2.64%, and 2.02%, respectively, which were 51.6%, 57.9%, 59.0%, and 71.9% lower than the original error rates, respectively. Among these, the DRO dataset has the greatest effect with errors dropped 3.5-fold. We found that the first iteration of error correction is usually the most effective. Although there was almost no effect after 3 iterations, we set 5 iterations as the default parameter value in our program to safeguard more difficult scenarios. Through an iterative error correction strategy guided by spatiotemporal information, ITEC improves greatly the accuracy of tracking and lineage reconstruction.

### Minimum-cost circulation model facilitates the error correction and helps maintain the integrity of tracks

While iterative error correction is conceptually straightforward, it is technically difficult to implement, due to the challenge in identifying errors or the concern of increased computational costs. We addressed both issues at the same time by adopting our recently developed computationally efficient and flexible model, the minimum-cost circulation model (*44*) (Fig. S5). Firstly, the model set a new record of computational complexity, which was further supported by the empirical evidence that it achieved an average increase in speed of 53 to 1192 times over three peer algorithms across multiple datasets (*44*). This improvement in efficiency relieved us from concerns about computation cost. Secondly, we introduced further modifications to the original minimum-cost circulation model to identify the localizations of the erroneous candidates for the error correction module. (1) To address the error of missing detections, frame-skipping connections were added (Fig. 3B and Fig. 3C). The model can tell whether a frame-skipping connection is turned on to fit the observed data. When the connection is turned on, it suggests there is a missing cell between the two connected cells. Then a module of detecting missing cells is called on to redetect them and splice originally broken tracks into continuous ones. (2) To tackle over-segmentation and under-segmentation errors, we add two arcs to the model and permit one-to-two and two-to-one matchings at specific time points (Fig. 3D and Fig. 3E). If the cost of splitting or merging cells is lower than that of the original segmentation result, and other biological or probabilistic constraints are satisfied, a decision is made to correct the segmentation result.

To enhance track integrity, the minimum-cost circulation model was further modified to allow for cell motion with large displacement. Due to individual cell differences and biological activities, some cells or cell populations have large motion within specific time periods (Fig. S8A). Therefore, we initially make a conservative estimate and only use high-confidence linkages to determine error corrections. Here, the tracking results obtained with the stringent linking standard are named tracklets. Eventually, large motion associations become increasingly reliable and more relaxed linkages can be tolerated. Finishing all the iterations of error correction, we abstract the existing tracklets into individual nodes (Fig. S8B) and apply the minimum-cost circulation model again to allow for a wider range of motions. This trick enables the implementation of error correction strategy and the merging of numerous broken tracks while ensuring high accuracy.

### ITEC enables long-term lineage reconstruction at terabyte level

Time-lapse 3D imaging datasets of embryonic development are often at terabyte-scale (*18*). In order to maintain accurate tracking during long periods of cell migration and division. Due to the accumulation of tracking errors, only high-fidelity tracking can lead to conclusions with practical biological significance. To our knowledge, current tracking methods often lack the capability to handle large-scale data, fail to output results in complex scenarios, have excessively long runtimes, or exhibit a high error rate during long-term tracking.

To test the capability of ITEC in long-term lineage reconstruction, we used a dataset consisting of 1001 time points (*39*). This dataset captures 16.7 hours of whole-embryo development in zebrafish, with a total data size of 1.5 TB (1001×253×1792×1818 voxels, T×Z×Y×X). We aligned the time window with the absolute developmental time, mapping it to the 5.5-11.3 hpf period (*49*). We ran our pipeline on this dataset in a distributed manner, dividing the 1001 frames into several batches with a certain overlap between adjacent batches. Subsequently, we associated the results from each batch using the nearest neighbor principle. To test the accuracy of these lineages, we randomly selected 31 cells at the first time point and then manually curated the corresponding lineages for the first 600 frames, which contained a total of 9.0 million cell instances with about 8,000 cells at the first time point and 21,000 cells at the end. Among over 30,000 linkage edges and a similar number of cell instances, a total of 91 errors were identified, which means the linkage accuracy of our method reached 99.7%. More notably, 15 lineages, that is, 48.4% of lineages, were error-free in the analysis spanning 600 frames (each lineage typically contains 1 to 3 divisions). Fig. 4A and Fig. 4B show the results obtained by ITEC.

To the best of our knowledge, it is the first time that the lineages of the whole-embryo containing up to 21 thousand cells at one time point and spanning 600 time frames can be fully reconstructed at the level of around 50% error-free and complete lineages. In contrast, the best peer methods are estimated to achieve only a 20% error-free rate. It is worth pointing out that as 10 seconds are usually needed to annotate one cell instance such an embryo with total 9 million cell instances would take 25,000 manual hours (more than 1,000 days) to be fully annotated. In contrast, ITEC can complete the same analysis in 8.7 days in a fully serial manner with a single processor and achieve results comparable to human labeling (Fig. 4D). If distributed operation is adopted, the analysis time will be shortened to 2-3 days.

### Capturing complex movements during embryonic development

Based on the long-term, high-fidelity tracking performance, we expected ITEC to be effective in capturing morphogenetic events. Analysis of the tracks during zebrafish embryogenesis (Fig. 4A) reveal epiboly movement, the movement of cells towards the vegetal pole, and convergent extension, the intercalary movement of cells towards the dorsal side (*50*). To further investigate whether ITEC is capable of capturing developmental dynamics at key stages, we analyzed internalization movements. Internalization is the defining cell motion observed during gastrulation. It is one of the most difficult motions for tracking, because cells ingress, undergo large-scale movements and rearrangements, and become less accessible for imaging (*50*). Starting from 50% epiboly at 5.5 hpf, the zebrafish blastoderm margin moves and folds inwards, generating an outer and inner layer, the epiblast and hypoblast, respectively (*51*). Cells in epiblast and hypoblast have opposite motion directions, move rapidly, and change neighbors, making cell tracking very challenging. Since involution takes place during early stages of embryogenesis, any tracking mistake at this stage carries forward to later stages, resulting in erroneous reconstructions.

Even with such a challenging dataset, ITEC captured internalization (Fig. 4C and Movie S3). We compared our approach with the classic algorithm TGMM (*1, 39*), which was especially designed for embryonic cell tracking. The most significant difference can be observed in the box in Fig. 4C at 8.3 hpf. Red cells represent involuted cells in the hypoblast, whose color indicates that their origins were from the blastoderm margin as tracked by ITEC. In contrast, TGMM cannot track these margin cells correctly and the internalization process was clearly missed (*52*). These results show that ITEC can reconstruct cell lineages over long distances and through rapid and complex cell movements.

### ITEC enables accurate fate mapping of multiple tissues at the single-embryo level

High-accuracy, long-term lineage reconstruction based on *in vivo* imaging provides a powerful approach for revealing complete fate maps, in which the earlier location of all the cells in the same embryo can be linked to its later position and fate. To test the potential of ITEC in fate mapping, we manually annotated the positions of several organs during zebrafish embryonic development at 11.3 hpf and then applied ITEC to trace back to their ancestors retrospectively to 5.5 hpf (Fig. 5A). The result demonstrated a high degree of consistency with existing fate maps reconstructed from *in vivo* clonal tracing methods (*53, 54*) (Fig. 5C). The positions of progenitors of the brain, eyes, and somites (future muscles) align well with the reported progenitor location in existing maps (*51, 54–57*). Another evidence that validates the correctness of ITEC is the accurate backtracking of somites. The four tracked somites (namely AA, AB, BA, and BB) are located in close proximity at 11.3 hpf. When mapped to 5.5 hpf, AA/AB and BA/BB are clearly assigned to the left and right hemispheres of the embryo without any mistake (Fig. 5B).

To further validate the capability of ITEC in fate mapping, we manually curated the lineages of 293 labeled descendants in somites at 11.3 hpf (Fig. 5D), where over 192 thousand cells were involved in total and 17 descendants at 11.3hpf were excluded due to poor imaging quality. For our curation process, we worked backwards from the end timepoint of the dataset, paying close attention to how critical events like cell division and large displacements were handled. On average, it took about 0.8 hours to curate a single cell’s entire 1001-time-point lineage, amounting to a total of roughly 220 hours for the entire curation. As a comparison, manually mapping all of the 276 descendants would require about 800 hours. After curation, we found that these somites originated from a total of 136 ancestors (Fig. 5E) at 5.5 hpf. Without curation, the automatic fate mapping of the 276 descendants identified 94 ancestors, of which 65% are the correct ancestors. Such a good fidelity enabled drawing biologically meaningful conclusion based on automatic fate mapping, which was confirmed in the next section. Compared to the conventional clonal analysis, ITEC enables the fate mapping of multiple tissues or organs on the same embryo, which empowers further analysis of the boundary forming of adjacent tissues and the relative developing time among different organs, as demonstrated in the next sections.

### ITEC reveals the dynamic formation process of boundaries from a mixed population of cells

Time-lapse imaging data, combined with an efficient tracking method, can reveal how intermingled cells segregate into defined structures and territories. Somites are the paired blocks of paraxial mesoderm that will become the adult muscles. During somitogenesis, the unsegmented paraxial mesoderm progressively segments into bilaterally symmetrical somites with defined boundaries (*58, 59*). To test the capability of ITEC in exploring cell intermingling and segregation, we analyzed somite formation. ITEC revealed the entire dynamic process of the formation of somites in the anterior part of the animal (Fig. 5B and Movie S5). The paraxial mesoderm of zebrafish derives from two symmetrical territories close to the margin that converge toward the margin, internalize and migrate along the anterior-posterior axis. Interestingly, ITEC allowed us to determine that at 5.5 hpf, progenitor cells that will give rise to separate somites exist in an intermixed state without discernible boundaries. In order to accurately quantify the formation process of the somite boundary, we fit the classification hyperplane to the somite pairs at each timepoint and defined the average misclassification ratio of the two classes as the mixing index (Fig. 5E). Starting from 6.7 hpf, the mixing index gradually decreases, indicating that in ipsilateral somites AB and BA, as well as AA and BB, cell populations become gradually separated, marking the initial formation of somite boundaries. We notice that although somites are bilaterial they are not completely symmetric. Somites AB and BA segregated earlier and faster than somites AA and BB. Until 9.0 hpf, the mixing index decreases to a very low level and fluctuates slightly, indicating that boundary structures are established as interfacial gaps emerge at their contact zones. Overall, starting at 6.7 hpf, the mixing index shows a monotonic downward trend, indicating the transition of somite progenitors from a mixed to a segregated state. Notably, measuring the mixing index without human curation led to the same conclusion.

We extended the analysis of the mixing index to brain and eyes (Fig. 5F). The ancestors of the brain and eyes were heavily intermingled at 5.5 hpf until having a more distinct boundary at 10.1 hpf, and significantly separated at 11.3 hpf, agreeing with previous reports (*54*). In addition, the left and right eyes exhibited a similar ipsilateral mixing pattern as the bilateral somites, with no contralateral intermixing from beginning to end. Given that the distance between the left and right eye progenitors was very close at 5.5 hpf, these results highlight the accuracy of ITEC in revealing the continuous spatiotemporal features of organ development and the gradual transition of cells from a mixed to a segregated state.

### Revealing the heterogeneity and distribution pattern of multiple regional origins

The developmental origin of midline structures – notochord, floor plate, spinal cord and more generally the body axis – and the diversity of cell behaviors that give rise to the embryonic midline have long been under debate (*60, 61*). Leveraging whole-embryo lineage tracing with ITEC, we revisited this classical problem by directly linking early progenitor positions to their later midline contributions.

We identified the midline of zebrafish embryo at 11.3 hpf and traced back the progenitor cells at 50% epiboly that contribute to the left and right hemispheres as well as to common bilateral progenitors. We found that these bilateral progenitor cells were symmetrically distributed along the midline formed by the shield, and originated from regions further away from the midline at the neural plate border (Fig. 6A). To analyze spatial asymmetries, we expanded the midline domain at 11.3 hpf outward to define three regions – the midline, left and right territories – and examined the distribution of their ancestral cells across the dorsal and ventral hemispheres. Cells originating from the dorsal hemisphere contributed proportionally more to the midline than those from the ventral hemisphere (Fig. 6A), and this asymmetry persisted as the midline domain expanded. This bias likely reflects the fact that dorsal cells approach the midline more directly through convergence movements during gastrulation.

Subsequently, we examined whether tissues and organs arise through the coherent migration of cell populations or through the reorganization of cells originating from distinct locations. To address this question, we selected cells and their neighboring populations at 11.3 hpf and traced their ancestors back to the 5.5 (early gastrula), 6.8 (mid gastrula), 8.4 (late gastrula), and 9.8 (bud) hpf stages, respectively. As shown in Fig. 6B, when a group of tightly arranged cells near the midline was selected, their ancestors were found to originate almost equally from the left and right hemispheres, supporting the hypothesis of cell reorganization. Between 5.5 hpf to 6.8 hpf, these ancestral cells underwent vegetal movement during epiboly and became increasingly dispersed on both sides of the embryo. Immediately afterward, during convergence, cells from both hemispheres aggregated toward the dorsal midline, ultimately forming the compact cell cluster observed at 11.3 hpf.

At 50% epiboly, the blueprint for the different germ layers of a zebrafish embryo is established (*50*), and secondary movements begin. We wondered if the different germ layers would show different speeds or cell dispersion capacities from the midline across gastrulation. We quantified these cell movement events using a mean distance dispersion index, which measures the degree of ancestral dispersion based on the average value of pairwise distances among the traced cells (Fig. 6C). The dispersion index of cells near the ectodermal midline was significantly higher than that of cells in other regions, indicating a broad and heterogeneous origin of cells near the midline. Notably, dispersion peaked near the end of epiboly, when cells were maximally distributed across the spherical embryo surface. Together, these analyses provide a novel perspective for verifying and quantitatively analyzing challenging embryonic development processes, including cell mixing and rearrangement, and enable the systematic revelation of the origin and overall behavior of cell populations across germ layers.

### Integration of ITEC with spatial transcriptomics identified correlation between cell dynamics and molecular programs

During embryonic development, the spatiotemporal expression of genes provides crucial insights into cell behaviors and fate decisions, but static measurements capture only snapshots at specific time points. In contrast, *in vivo* imaging reveals dynamic processes such as cell migration and differentiation, yet lacks comprehensive molecular information about the underlying gene expression programs. To bridge these complementary approaches, we applied our recently developed technique, weMERFISH, a whole-embryo spatial transcriptomics platform based on multiplexed error-robust fluorescent in-situ hybridization (*9*), which captures gene expression patterns across the entire embryo. In our previous work, we developed MERFISH-FATE, which integrated whole-embryo spatial transcriptomics with *in vivo* cell tracking to reveal how gene expression patterns evolve through morphogenetic movements during gastrulation, and showed that similar spatial patterns can arise through distinct transcriptional dynamics. Building on this framework, we characterized in more details the correlation of gene expression with multiple morphodynamic features. We integrated the spatial coordinates of weMERFISH data with ITEC by applying a globally optimal matching algorithm (Fig. 7A) to align corresponding regions between the 50% epiboly and 75% epiboly stages. Because of embryo-to-embryo variability, this alignment represents an approximate correspondence of tissue regions rather than single-cell matches, enabling the exploration of how gene expression dynamics relate to morphogenetic cell movements at the tissue and organ levels.

We used the lineage reconstruction results to calculate a cell migration velocity map at the 75% epiboly stage (Fig. 7F). Cells located at the margin exhibited the highest migration speeds, with dorsal cells showing the fastest movements consistent with convergence during gastrulation. To link cell dynamics with molecular programs, we computed the correlation coefficients between the local migration velocity and gene expression levels (Fig. S10). Fig. 7G shows the top ten most correlated genes, which include *itgb5, nradd, fgf24, wnt8a*.*1*, and *wnt5b*. We then performed Gene Ontology (GO) enrichment analysis on genes with the strongest correlations (Fig. 7H). The most significantly enriched categories included somite and tail development, as well as components of Wnt and Planar Cell Polarity (PCP) (*wnt8a, lef1, wnt5b, wnt11f2, ptk7a*) (*62–64*), Fgf (*fgf8a, fgf24*) (*65*) and Notch (*dld*) (*66*) signaling pathways, suggesting that these signaling cascades might be linked to the characteristic fast motion of cells during convergent extension.

The local coherence of cell velocities across different regions reflects the tissue mechanical properties such as fluidity index (*67, 68*). To relate these properties to gene expression, we calculated the local variance of velocity within neighborhood of 50 cells (Fig. S12) and identified genes whose spatial expression patterns resembled the variance map (Fig. S11). We found that the spatial heterogeneity in cell movement is associated with developmental programs that control mesodermal remodeling, vascular morphogenesis, and axis patterning, such as genes *hmcn2*.*1, nid2a* and *tcf7*, highlighting increased fluidity in the mesendodermal tissues (*69*) and the molecular basis of mechanical and behavioral diversity across the embryo.

The integration of ITEC with spatial transcriptomics enables direct connections to be drawn between gene expression patterns and cell dynamics, helping establish a solid foundation to elucidate the spatiotemporal mechanisms underlying cell behavior and providing a new perspective for studying developmental processes.

### Molecularly defined regions are asynchronously formed

Using the cell mapping relationships derived from live cell tracking, we projected the cell-type identities at the 75% epiboly back onto the 50% epiboly stage (Fig. 7B) to investigate the mechanisms underlying boundary formation between transcription-defined regions. This process is similar to fate mapping, with the key difference lying in the source of the region annotations, which were obtained through weMERFISH spatial transcriptomics (*9*). Among the 14 regions with class labels, we analyzed the mixing at the boundaries of 22 region pairs, excluding non-adjacent region pairs and those belonging to the different layers. We fitted a separating hyperplane for each pair of regions and defined a mixing index as the sum of the proportions of misclassified cells on both sides of the hyperplane.

Through clustering the distribution of the mixing index at the two epiboly stages, three distinct mixing patterns emerged (Fig. 7E). One example for each category was shown in Fig. 7D and more examples were detailed in the supplementary figures Fig. S13–S15. We found that 9 region pairs maintained relatively clear boundaries at both the 50% and 75% epiboly stages (Fig. S13). For example, somitic mesoderm and notochord clusters maintained clear boundaries at both stages, consistent with the conclusions we previously obtained (*9*) (Fig. 7D). Additionally, 7 region pairs exhibited strong mixing at 50% epiboly, but the intermixed cells segregated to molecularly and anatomically distinct clusters at 75% epiboly (Fig. S14), similar to the patterns we observed in somite segment formation at much later stage of 11.3 hpf (Fig. 5F). For example, lateral anterior mesoderm and somitic mesoderm cells were much intermingled at 50% epiboly but became completely segregated at 75% epiboly (Fig. 7D). Furthermore, 6 region pairs remained on a relatively mixed level at 50% through 75% epiboly stages (Fig. S15). Note that the non-separated boundaries for these region pairs at 75% epiboly may be partly due to the inaccuracy from the registration between individual embryos.

We were also interested in the directionality of mixing at the boundaries, so we further analyzed the directionality of cell mixing across all regions by quantifying the cell mixing indices along the anterior-posterior (AP) and medial-lateral (ML) axes, respectively (Fig. 7C). This analysis showed that the cell mixing along the AP direction was significantly greater than that along the ML axis (Wilcoxon test, p=4.428×10^−4^). All these results demonstrated that the molecularly and anatomically defined regions were asynchronized and formed at different epiboly stages, likely reflecting different underlying mechanisms.

## Discussion

Accurate lineage reconstruction in 3D time-lapse embryonic development dataset has long been a major challenge in the field of developmental biology. Due to factors such as the heterogeneity of image quality, the vast scale of data, and the extremely high requirements for accuracy, current mainstream methods have not yet truly solved this problem. To address these challenges, we propose ITEC, a cutting-edge pipeline designed for embryo tracking in live-imaging microscopy dataset, with the ultimate goal of reconstructing complete and accurate lineages. We fully exploit the spatiotemporal information contained in the multi-dimensional data, and skillfully utilize our novel error correction strategies. This enables mutual promotion between cell detection and linkage, forming a stable closed-loop feedback system, thereby significantly enhancing the accuracy and robustness.

ITEC has achieved a significant lead in tracking accuracy compared to the state-of-the-art methods. We validate our approach on embryonic development datasets of common model organisms such as zebrafish, mouse, and *Drosophila*, and ITEC consistently yields the best performance—with an average error rate reduction of 58.5% compared to the second-ranked method. Our method also achieves the highest lineage continuity and overall accuracy in the evaluation of long-term lineages. The lineage reconstruction task imposes extremely high standard on accuracy: a single error among hundreds or thousands of frames can lead to mistakes in cell lineage tracing and fate mapping, thereby rendering the results unreliable. We believe that ITEC, with an unprecedentedly low error rate, has made this once-impossible task feasible. It is expected to facilitate the emergence of a new paradigm in developmental biology research of both model organisms and *in vitro* models such as organoids and embryo-like structures.

Then, what exactly enables ITEC to achieve such high accuracy? We attribute this to the full utilization of spatiotemporal information. While other methods also leverage temporal consistency to a certain extent, our method takes the thorough exploration and refined the feedback of this information. By analyzing and examining numerous cases where the tracking errors occur, we have designed a comprehensive error correction module. This module, when integrated with a minimum-cost circulation framework, identifies potential errors from the initial tracking results, corrects them, and achieves stability through multiple iterations.

We validated the performance of our method on data spanning 1,001 frames and terabyte scale. Furthermore, we utilized these results to conduct analyses of fate mapping and the findings are consistent with existing advancements in developmental biology, verifying the accuracy of our method. Previous research used clonal analysis by single-cell transplantation or fluorescent labeling to analyze cell fate by a sum of several embryos (*51, 54–57*). However powerful, this method can miss fuzzy tissue boundaries or will miss the fate and lineage relationships of the unlabeled neighbors. Using ITEC, we could investigate the formation process of zebrafish somite development from a dynamic perspective and explored the origin of midline structures *in toto*, providing biologists with a novel method that combines cell-level tracking with organ-level research. Additionally, we quantified a series of important kinetic parameters during embryonic development, such as the dispersion index of cell origin, cell migration speed, and variance from a quantitative viewpoint. We also integrated *in vivo* imaging with spatial transcriptomics, and based on this, explored issues including organ boundary formation and the relationship between cell migration and gene expression. These studies strongly demonstrate that our method not only achieves excellent performance in terms of tracking accuracy but also holds great potential for advancing biological research.

To provide convenience to the broader community, ITEC strives to be user-friendly. Therefore, during development, we retained parameters with biological or image-related significance for users to adjust, and concretized complex principles into several key parameters. In addition, ITEC allows users to adjust parameters via a visual interface and integrates tracking results with Mastodon (*47*), a large-scale cell tracking platform, facilitating users’ downstream analysis.

Despite its advantages, ITEC still has some limitations. First, ITEC’s runtime can be further optimized: for large-scale datasets, if users do not adopt a distributed computing strategy or have no access to parallel computing solutions, the analysis time will be relatively long, which may affect the user experience. Going forward, we will improve operational efficiency and user experience by designing better parallel strategies and optimizing the underlying time-consuming modules.

Additionally, our method was originally designed for the identification and tracking of nucleus-labeled embryonic cells. It would be suboptimal when segmenting cells with diverse morphologies—such as polyhedral, columnar, or spindle-shaped cells—or cells with heavily uneven internal expression. Although certain corrections can be achieved through the error correction module, we aim to expand the versatility of the ITEC framework in future work, thereby enabling the segmentation and tracking of other cell types.

In the future, we also aim to integrate embryo tracking with other biological technologies (*9, 70–72*). For instance, while we currently use nuclear channels for tracking, integrating with membrane channels is expected to enable the exploration of morphological dynamics. Additionally, the deeper integration of embryo tracking and spatial transcriptomics on the same embryo is expected to reveal the correlation between developmental rhythms and gene expression, thereby advancing our understanding of embryonic development from multiple perspectives.

## Acknowledgments

We thank Li Yu, Jianquan Ni, Cuifang Zhang, Aibin He for discussions and assistance with the project; Donglin Li, Jian Zhong, and Jianqiong Hu for their contributions to the experiments; Philipp Keller and Kate McDole for providing the embryo development dataset. We thank the Biozentrum Imaging Core Facility at the University of Basel for technical support. We acknowledge the support of Imaging Core Facility, Technology Center for Protein Sciences, Tsinghua University for providing instrumentation and technical assistance from Wenjuan Wang, Yanli Zhang, Dan Zhang, and Yuke Feng. We thank Alba Aparicio Fernandez, Rita Gonzalez and Diana Medeiros Gomes for zebrafish husbandry. We thank the Yu lab and the Schier lab for discussion.

## Author contributions

G.Y., Y.W., J.N.A., and A.F.S. conceived the research. C.W. designed the first version of ITEC and M.W., Q.Z, W.Z. developed the pipeline. M.W., Q.Z. and Y.C. performed the experimental evaluation. Y.W., J.N.A. provided the zebrafish embryo data and analyzed the biological results with supervision from A.F.S., Q.Z., M.W., Z.M., X. C., Y.C., W.Z., B.Y. analyzed the results and created the figures. Q.Z., M.W., J.N.A., Y.W. wrote the manuscript. G.Y. provided guidance. All authors contributed to editing the paper.

## Data availability

- The tracking evaluation results of ITEC and peer methods in this paper, as well as the ground truth we used are shared through Mendeley with the dataset identifier https://dx.doi.org/10.17632/tg55phtk4r.1.
- The FISH1 (*9*) and FISH2 (*41*) data will be shared by the lead contact upon request due to the large amount of data.
- The MOUSE (*1*) data can be downloaded fromhttps://idr.openmicroscopy.org/webclient/?show=project-502.
- The weMERFISH (*9*) website: https://schier.merfisheyes.com/.

## Code availability

All original code has been deposited at https://github.com/yu-lab-vt/ITEC.

## Supplementary Materials

### Live imaging datasets

All zebrafish experiments were performed according to the Swiss Law and the Kantonales Veterinäramt of Kanton Basel-Stadt (licenses #1035H). The FISH1 dataset was produced in the Schier lab in the University of Basel (*9*). The dataset recorded the early stage of the development process of a zebrafish embryo. Staging of the embryos was performed according to (*49*), available at the Zebrafish Information Network (ZFIN) (*73*). There are more than one million cells in 192 time points. We used our method to generate a preliminary tracking result first, and then a part of the tracks with biological interests were selected and professionally curated twice. As a result, we generated 37 lineages containing more than 32 thousand associations in total.

The FISH2 dataset is from Keller lab in HHMI Janelia Research Campus, and has been demonstrated in (*39*). It recorded a zebrafish embryo as well but in a relatively longer time duration. We selected 100 time points (300-399) at the middle stage as a complement of FISH1, which had the same amount of cells. We also manually curated based on our results and selected 36 lineages as the ground truth.

The MOUSE dataset is from Keller lab in HHMI Janelia Research Campus recording the development of mouse embryo. Besides, it serves as the evaluation dataset in Linajea (*41, 42*). We selected the middle stage with 50 frames (250-299) for evaluation, which offers a moderate level of difficulty. Due to occasional errors and ambiguities in the annotations, we combined publicly available annotations with our own corrections and annotations to create the final evaluation ground truth.

The DRO dataset is from Hufnagel group in EMBL Heidelberg (*74*) recording the development of *Drosophila melanogaster* embryo, and has been partly annotated and tested in (*75*). The annotation is claimed as a combination of automatic segmentation with refinement and manual association. However, we noticed the existing of mistakes, such as broken tracks. We manually checked some annotations and selected 62 correct lineages as the ground truth to ensure reliable evaluation.

### Peer methods

The first one is Tracking with Gaussian Mixture Models (TGMM) (*1, 39*), which is an unsupervised algorithm that uses watershed (*76*) to generate segmentation candidates, and then uses 3D Gaussian mixture model to associate cells in two adjacent time points greedily and determine the final segmentation, where results in every time point are also the prior for the next time point.

The second one is Linajea (*41, 42*), a two-stage algorithm whose detection is based on 3D U-Net (*77*). The tracking stage considers all cell associations in the whole data including divisions, formulates them as an integer linear programming (ILP) problem, and can obtain a global optimum theoretically. It is unavoidable to bring NP-hardness as mentioned before, while the time and memory consumptions are unacceptable for large-scale data. As a result, Linajea processes real data block-wise and is content with an approximate solution to the original optimization problem.

The third one is Ultrack (*38, 43*), which handles the detection and tracking task jointly. It uses Ultrametric Contour Maps (UCM) (*78*) to generate segmentation candidates hierarchically, and then formulates the tracking problem into ILP similar to Linajea. Ultrack does not determine the segmentation results at the beginning. Instead, it selects the optimal segmentation results from hierarchies while solving for the tracking results.

### Evaluation metrics

Due to the sparse labeling of cells in ground truth, we define that a detected cell is successfully recognized if it is the closest one to a cell in the ground truth, and the distance is smaller than the cell radius (Fig.S1A). A detected cell can be matched to at most one ground truth cell and vice versa.

#### Detection FP/FN

If the centroids in the densely annotated ground truth are less than a certain distance from the centroids in the detection results, we match them and calculate FP/FN based on this (Fig.S1B).

#### Euclidean distance metric

For each annotated centroid, we determine which nucleus it was assigned to in the automatic reconstruction. Then, we measure the Euclidean distance between the annotated centroid and the centroid of the nucleus from the automatic reconstruction. If the detection results have severe under segmentation, this metric will be biased higher (Fig.S1C).

#### Nearest neighbor normalized distance metric

For each annotated data point, for which we have calculated the Euclidean distance metric explained above, we normalize the distance measure by the distance to the nearest neighbor in the automatic reconstruction. If there is severe over segmentation in the detection results, the normalized metric will be biased towards a larger value (Fig.S1D).

#### Linkage TP/FP/FN

For each cross-frame association, an association is regarded as a true positive (TP) if both the parent and daughter cells are successfully recognized and match an association in the ground truth. If at least one cell is recognized but the association is wrong, it is treated as a false positive (FP). If an association in the ground truth cannot find any matched cell or if the matched cells don’t have a track, the association is miss detected and treated as a false negative (FN) (Fig.S1E). Since cell division is a process that occurs over a period of time, we adopt a lenient evaluation strategy for divisions that occur earlier or later than in the ground truth, meaning they are not counted as FP or FN errors (Fig.S1F).

#### Errors per ground truth edge

We define FP/FN/Sum per ground truth edge as the ratio of FP/FN/(FP + FN) to the number of edges in the ground truth, respectively. These three metrics can quantitatively evaluate the performance of tracking.

#### Average error-free length

We use the complete lineage annotated from beginning to end as the unit, average error-free length is the number of consecutive time points over which we encountered no tracking errors. Due to imaging defects, some lineages in the ground truth are incomplete, so we weight them by track length (Fig.S1G).

#### Proportion of error-free tracks

This metric represents the proportion of completely correct lineages to the total annotated lineages (Fig.S1G).

### Application scenarios

#### Low signal-to-noise ratio

ITEC accurately models the noise and has several feedback ideas to iteratively improve tracking accuracy. Therefore, it has an absolute advantage in processing low signal-to-noise ratio data, which is difficult to achieve based on deep learning methods.

#### Large scale data

In segmentation module, ITEC adopt a parallel strategy to accelerate seed growth and boundary generation. In tracking module, it applies the most advanced machine learning techniques for data association, and despite multiple iterations, it can still achieve high running speed.

#### High accuracy requirement

ITEC achieves high tracking accuracy through effective error correction module design. This is crucial in the reconstruction of the entire embryonic developmental lineages. ITEC can almost achieve zero error rate in the parts with clearer imaging, ensuring the integrity and accuracy of the lineages.

#### Multi application scenarios

ITEC has been applied in various embryos such as zebrafish, mouse, *Drosophila*, etc., achieving the highest performance. Meanwhile, it can handle both local regions of interest and the entire embryonic development process, with a wide range of application scenarios and potential.

### User-friendly design

ITEC strives to be user-friendly, primarily manifested in three aspects. First, it is fully based on unsupervised machine learning techniques, which eliminates the need for data annotation. Users only need to adjust parameters that have practical biological significance or are intuitively adjustable (Table S3). For example, we set thresholds with clear meanings for grayscale values, such as *background intensity, intensity upper bound*, and *intensity lower bound*. Users can easily obtain these parameters using mainstream image processing software available today. Additionally, there are parameters for balancing false positives (FP) and false negatives (FN), including *curvature threshold* and *division threshold*, which allow users to control the number of detected cells and confidence level according to their specific needs.

Second, we simultaneously design a UI interface and provide a parameter table, enabling users to conveniently adjust parameters and run the pipeline across different devices. Meanwhile, we have integrated the tracking results with Mastodon (*47*), a mainstream large-scale embryonic cell tracking Fiji plugin (*48*). This allows users to easily use the platform to view tracking results, lineage trees, and other information, as well as curate and analyze the results.

Third, we provide users with a comprehensive user manual, which includes multiple example parameter tables tested during our development process. This facilitates quick user onboarding and enables users to rapidly adjust parameters in practical applications.

### Distribution dispersion measurement

The mean distance dispersion index reflects the overall dispersion degree by calculating the average distance between all pairs of points. A higher value of this index indicates a more dispersed cell distribution. This index can well take the contribution of all cells into account and avoid being affected by outlier points. We define the coordinates of the points as *x*_*i*_ (*i* = 1, 2, …, *n*), and their average distance dispersion index is:

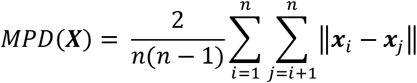

### Velocity map and variance map calculation

At the TM230 of FISH2 (corresponding to 75% epiboly), the velocity of each cell was derived from the cell tracking results of the 20 preceding and subsequent frames. Subsequently, the velocity of each cell position was obtained by averaging the velocities of the 10 neighboring cells around that cell. The variance of each cell is calculated from the velocity variances of up to 50 cells within a certain range around that cell.

## Online methods

### Step 1: Pre-processing

#### Deconvolution

In general, the output of an optical imaging system can be described as a convolution between the input signal and the point spread function (PSF) with additive noise *O* = *h* ∗ *I* + *N*, or in the frequency domain,

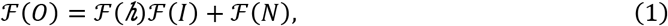

where *I, O, N* are the input time-series signals, output videos, and noise. *h* is the time-invariant PSF and *F* is the Fourier transform. Especially for fluorescence microscopy, deconvolution is crucial to restore the real signal I and can help to avoid under-segmentation. We formulate the deconvolution problem as a least squares estimation when *h* is given:

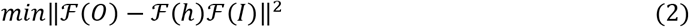

Rather than deriving the solution ℱ(*I*) = [ℱ(*h*) ∗ ℱ(*h*)]ℱ(*h*) ∗ ℱ(*O*) directly, to solve the ill-posed problems and suppress the noise, we use the Landweber algorithm (*79*) to estimate ℱ(*I*) iteratively:

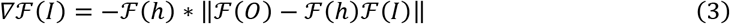

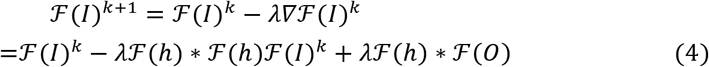

where *λ* is the step size and set to 0.9 in all experiments. If *h* is unknown, as a rule of thumb, an appropriate Gaussian kernel along the z-axis performs better than blind deconvolution approaches.

#### Rigid registration

Similar to most cell tracking methods, ITEC assumes the cell motions are small, which may not always be true for real data. Cell motions can be composed of three components: global motion mainly because of the whole embryo motion and rotation during experiments, tissue-level motion because of the development, and cell-level nearly Brownian motion. To eliminate the influence of global motion, we get the rigid transformation between adjacent time points *I*(*t*) and *I*(*t* + 1) by mean squared error minimization:

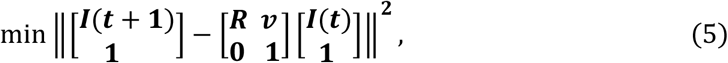

where *R* and *v* represent the rotation matrix and motion vector. It is worth mentioning that we do not transform the image data but only record the transformation matrix. Transformed data will consume much larger storage but lose more information than the original data. Rigid registration is also applied to stitch different angles of the same time point, which will be discussed in the post-processing.

#### Motion flow estimation

Rigid registration can make most cells satisfy the small motion assumption, while exceptions still exist. We treat these motions as tissue-level and applied patch-wise and pyramidal Lucas-Kanade optical flow (*45*) to roughly estimate these motions and register images non-rigidly.

Optical flow is a broadly applied motion estimation approach. However, the dense optical flow has quite expensive computation and storage overhead while the sparse optical flow is not applicable since cell locations may change during the error correction stage. As a compromise, we apply patch-wise and pyramidal Lucas-Kanade optical flow to roughly capture cell motion and register images non-rigidly. The basic idea is to downsample images and roughly estimate a global motion as initialization, and then upsample images, divide them as patches, and estimate the motion of every patch independently. The upsampling and patch-wise motion estimation can be executed multiple times while maintaining the same patch size but a smaller field of view for a patch.

To start with, we introduce the method of rigid registration through optical flow under the following assumptions: (i) The cell motion is small. (ii) The cell intensity does not change too much in two images. (iii) Adjacent cells share similar motions. The small motion assumption will be satisfied when applying pyramidal optical flow, and the other two assumptions are naturally satisfied by embryonic imaging data. Based on the assumptions, for a pixel at location (*x, y, z, t*) in a video *I*, it should have the same intensity at the next time point:

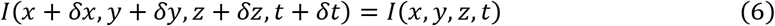

where (*δx, δy, δz*) is the motion distance along coordinate axes. Because of the small motion assumption, we take the first-order approximation:

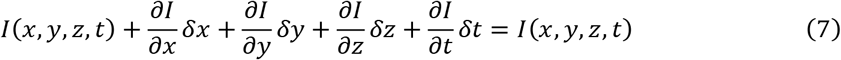

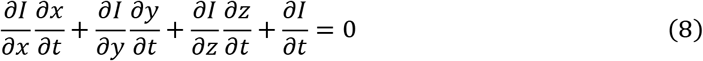

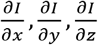 are image intensity gradients along different directions, and 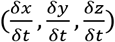 is the cell motion velocity waiting for estimation. To robustly estimate the motion velocity, we consider *n* multiple pixels (*x*_*1*_, *y*_*1*_, *z*_*1*_, *t*), (*x*_*2*_, *y*_*2*_, *z*_*2*_, *t*), …, (*x*_*n*_, *y*_*n*_, *z*_*n*_, *t*) in one image or a patch. Notably, considering all pixels may not bring better estimation since some background pixels are pure noise. Then we have

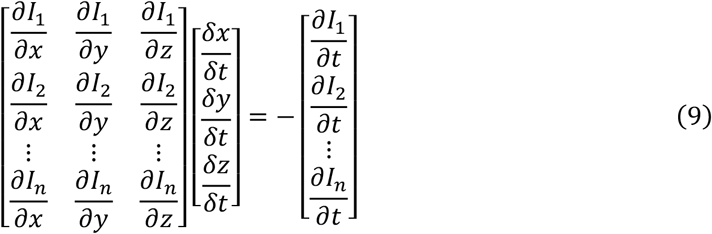

or in short,

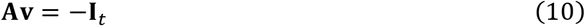

Then 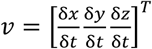 *c*an be solved by least squares with the given objective function:

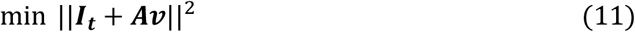

The optimum **v**^∗^ = −(**A**^*T*^**A**)^−1^**A**^*T*^**I**_*t*_ is the estimated velocity. Then we upsample the image, divide the image as patches, and refine the velocity estimation with *x*\* as an initialization. The following estimations are exactly the same as the first one if the patches are processed independently. However, some patches may be at the embryo boundary, contain very few foreground pixels, and cannot be correctly estimated without the help of neighbors. It is natural to assume adjacent patches should have similar motions and consider the similarities when estimation. We assume that there are m patches, describe the patch relationship as a graph, and use the mean squared error to describe the similarity cost:

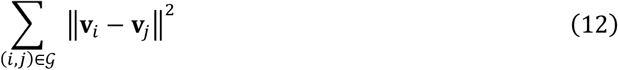

Finally, the objective function is the least squares estimation of patch motions with the regularization of similarities:

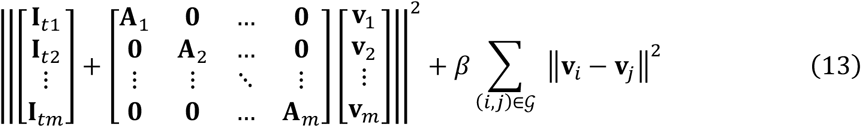

where *β* a regularization factor. In general, we prefer to set *β* to 1~5% of the mean intensity, which can contribute but not dominate the estimation. The optimal solution 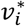 will be treated as the motion velocity of the center of patch *i*. Therefore, for any location in the image, we can estimate its velocity through the interpolation of adjacent patch centers.

### Step 2: Initial Detection

Accurately identifying cells within complex, crowded populations and performing pixel-level segmentation presents significant challenges. On one hand, strict differentiation between noise and genuine signals is required. Without precise noise modeling, many noise artifacts can be misclassified as cells. On the other hand, intrinsic imaging heterogeneity leads to intercellular variations. While most cells appear bright in the center and dark at the periphery, some exhibit internal textures resembling multiple cells, potentially causing over-segmentation or under-segmentation. To address these issues, first, we perform robust estimation of background noise by assuming noise follows an independent Gaussian distribution and estimating its variance to establish a foundation for subsequent steps. Second, because the maximum principal curvature (short for principal curvature or curvature in this section) encapsulates information about cell boundaries and intercellular gaps, we compute it and integrate results from the first step to accurately model and quantify the curvature distribution of noise (*46*). This separates cells from the background, excludes clearly non-cellular regions, and provides coarse positional information. Third, we utilize curvature information to grow cell boundaries, achieving precise pixel-level segmentation for enhanced downstream tracking (Fig. S4).

#### Variance estimation

To model background noise, we first require accurate estimation of its variance. We employ a robust variance estimation method assuming independent Gaussian noise in the background. This involves applying small-scale filtering to the image, then estimating the true variance using the distribution of intensity differences between each pixel and its neighbors, with correction coefficients generated via simulation. This approach leverages local information to mitigate global intensity variations under unknown signal distributions. The operation is performed on each z-slice of the 3D image, with the median variance across z-slices serving as the final estimate.

#### Boundary map generation

Principal curvature provides essential characterization of cell boundaries, particularly in dense populations, where it prevents over-merging and improves segmentation accuracy. However, curvature is sensitive to intracellular noise and struggles to distinguish internal textures from noise, making it necessary to perform smoothing. With small smoothing parameters, cell contours are preserved clearly without distortion, but cells with gaps may be over-segmented, making this suitable for boundary extraction. With large smoothing parameters, intracellular regions become smoother and more homogeneous, making this suitable for cell identification.

To fully leverage this information, we implement a multi-scale principal curvature (MSPC) method based on order statistics through computing the normalized principal curvature with specific smoothing factors derived from the maximum eigenvalue of the modified Hessian matrix.

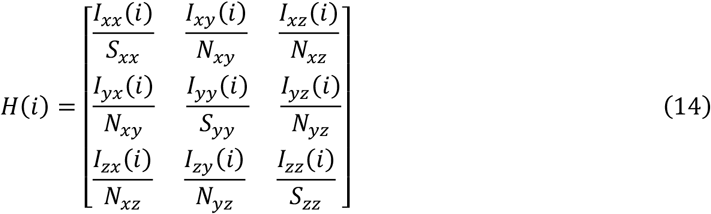

where *I*_*xx*_, *I*_*xy*_, *I*_*xz*_,… are the second-order partial derivatives of the smoothed image evaluated at the pixel *i*; *S*_*xx*_, *S*_*yy*_, *S*_*zz*_ is the sum of all pixel’s squared value in the second-order derivative of the Gaussian smooth kernel along the specific direction; The *N*_*xy*_ is the 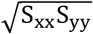.

We introduce two key enhancements. First, due to the z-direction characteristics of our data, we compute both 3D curvature and 2D curvature for each z-slice. Second, using the previously estimated background variance, we improve the curvature normalization across filtering scales. For 2D and 3D, background curvature follows a Gaussian distribution, so we normalize curvature at all scales to a standard normal distribution. Our current objective is to select cellular core regions. Based on accurate noise modeling and after excluding implausible filtering parameters, we use the minimum principal curvature across all scales to indicate seed information, applying a tunable threshold (typically −5) where values below this denote seeds. Simultaneously, we use the maximum principal curvature across all scales to indicate boundary information, where values above +3 are classified as boundaries and excluded from seed regions. This approach fully leverages the characteristics of principal curvature across different filtering scales, yielding more robust and accurate results.

#### Min-cut seed growth

The boundary growth is used to grow the seeds obtained in the previous section to obtain accurate cell boundaries. In fact, the growth of cell boundaries is to find the continuous pixels around the seed with the maximum sum of principal curvatures. Therefore, we model the problem as a minimum cut optimization problem with one source and multiple sinks.

For a specific seed *n*, we define the pixels belonging to the cell region grown from it as *R*_*n*_, and all other pixels as 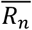. The boundary between these two groups can be represented using pairs of pixels, *C*_*n*_. Finding the set of boundary pairs *C*_*n*_ that minimizes the sum of *G*(*i*) and *G*(*j*) across all pairs:

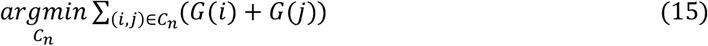

Here, 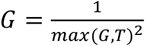, where *G* is the map of principal curvature for each pixel, and *T* is a minimum capacity threshold introduced to enhance robustness. It is worth noting that we combined the curvature of 3D and 2D to calculate *c*_*ij*_, in order to compensate for the erroneous growth caused by missing *z*-direction information. In this way, the problem of finding the boundary pixels with the maximum value in the curvature map *G* is transformed into finding the minimum values within *G*. This objective function also indirectly minimizes the boundary length.

To efficiently solve this problem, the problem is reformulated as a minimum cut problem. As shown in Fig. S4, the optimization graph constructed for Seed 1 includes all foreground pixels, with each node *i* representing a pixel. An edge *e*_*ij*_ connects every pair of neighboring nodes, and its capacity is defined as *weight*(*e*_*ij*_) = *G*(*i*) + *G*(*j*). All pixels corresponding to the seed are considered the source, while the background and other seeds are considered the sink. With this graph design, the minimum cut between the source and sink is calculated. This minimum cut represents the cell boundary, and the pixels enclosed by it are labeled as the detection result for the corresponding seed.

### Step 3: Tracking

#### Problem statement

The problem of object tracking can be formulated as an MAP problem. Here we discuss the most widely used formulations that consider only one-to-one matchings (*80*).

Let 𝒳 = {*x*_*i*_} be a set of detections, where *x*_*i*_ is a vector containing the position, appearance, and time index of detection *i*. A track 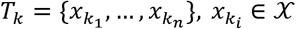 is a ordered list of detections and

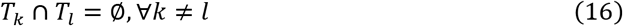

An association hypothesis is a set of non-overlap tracks 𝒯 = {*T*_*k*_}. Note that the 𝒯may not cover all the detections of 𝒳 as detections can be false positive. The purpose of identity inference is to find the hypothesis 𝒯 with the highest posterior probability:

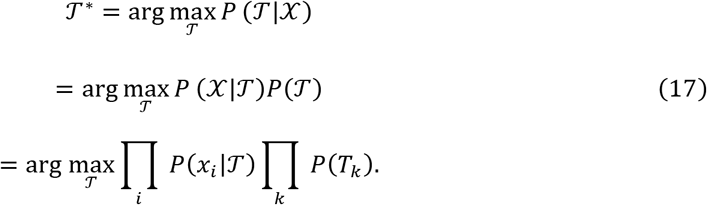

assuming conditional independence of detections given the hypothesis 𝒯 and independence between tracks, i.e. objects move independently.

Assume each *P*(*x*_*i*_|𝒯) follows a unique Bernoulli distribution *B*(1, *θ*_*i*_), with a preset parameter *θ*_*i*_ indicating the probability that *x*_*i*_ is mistakenly detected and thus should be excluded in the tracks:

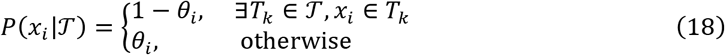

Since we only consider unary and pairwise relationships between detections, a track 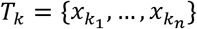 can be modeled as a Markov chain whose probability is

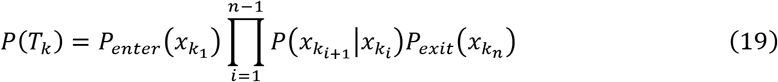

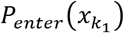 is the probability that 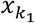 is the initial point of track *T*_*k*_. Similarly, 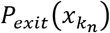 is the probability that 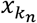 is the terminate point of track *T*_*k*_. By taking the negative logarithm of all the probabilities, and remapping 𝒯 into indicators 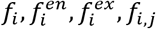, the MAP problem can be converted to:

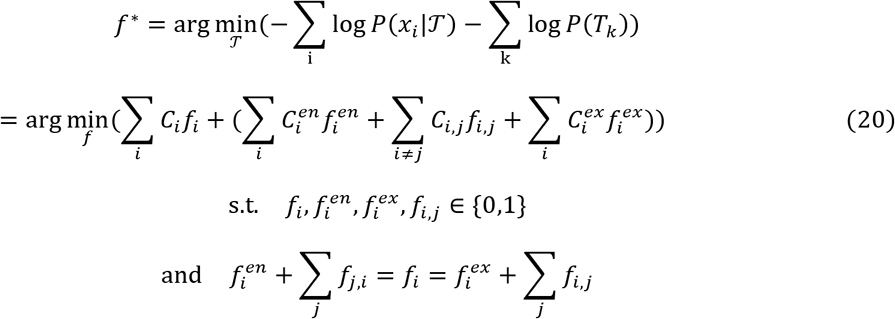

with

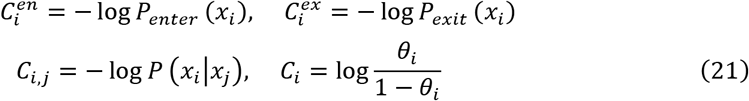

Here *f*_*i*_ = 1 indicates that detection *x*_*i*_ is included in a track of 𝒯 and *f*_*i*_ = 0 otherwise. *P*(*x*_*i*_ |𝒯) can be rewritten a 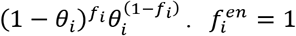 or 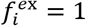 indicates that *x*_*i*_ is the initial or terminate point of a track in 𝒯. *f*_*i,j*_ = 1 means that detection *x*_*i*_ is followed by detection *x*_*j*_ in the same track in 𝒯.

Constraints in Equation 20 indicate that each detection can participate in at most one track and there is no splitting or merging of any track. We can see *f*_*i*_ = 1 forces detection *x*_*i*_ to be incident with at most one previous detection and at most one following detection in a track, so *x*_*i*_ will participate one and only one track. On the other hand,*f*_*i*_ = 0 automatically rules out the detection *x*_*i*_ from participating in any track, where 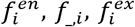, and *f*_*i*,__ will all be zeros.

#### Calculate affinity score between cells

A key component for a good tracker is the affinity score measuring the similarity among detections. However, for objects with a flat appearance like cells or particles, discriminative features are hard to design. From our observation, the motion and morphological changes of cells are relatively slow across time. Thus, we proposed to design an affinity score on the basis of the following assumption.

##### Assumption 1. Reasonably high temporal resolution of imaging

The temporal resolution is high enough such that, between any two consecutive frames, a cell in frame *t*+1 is the spatially closest one to itself in frame *t* among all cells in frame *t*+1.

Such an assumption is inevitable considering the lack of texture and morphology differences among cells. Based on this assumption, the most effective and faithful affinity score design is to measure the morphological similarity like the territory overlapping ratio between two detections.

However, either overlapping ratio or center distance suffers from the ellipsoid shape of cells. As is shown in Fig. S6, the two detections in (a) have the same overlapping ratio with those in (c), while has the same center distance as those in (b). However, from our point of view, the cells in (a) have smaller morphological changes than (c), but the evidence is not as strong the those in (b). Thus the desired score be able to differentiate these three conditions. As a single standard cannot achieve this task, we proposed the new design jointly consider these two criteria.

Let’s use *D*_*i*_ and *D*_*j*_ to denote the two adjacent detections as is shown in Fig. S6. *p* ∈ *D*_*i*_ represents a pixel *p* located inside*D*_*i*_. For two pixels *p* ∈ *D*_*i*_ and *q* ∈ *D*_j_, we define their distance *d*(*p, q*) as their Euclidean distance. We first define the distance from a pixel *p* to a detection *D*_*j*_ as

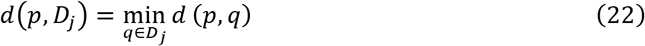

Then we can define the distance from detection *D*_*i*_ to *D*_*j*_ as

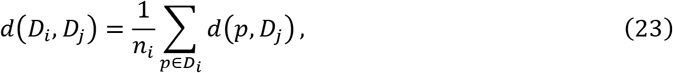

where n_i_ is the number of pixels in D_i_. With the score in Equation 23, if detection D_i_ has larger ratio of pixels overlapped with D_j_, these pixels will contribute zero to the cumulative distance, but weight down the average distance. Thus the two conditions in Fig. S6A and Fig. S6B will have different distances. For Fig. S6A and Fig. S6C, the non-overlapping pixels in D_i_ in Fig. S6A clearly has smaller average distance to D_j_ than those in Fig. S6C and thus can also be differentiated. The distance in Equation 23 is directional, which means d(D_i_, D_j_) ≠ d(D_j_, D_i_) holds for most of the cases. If we want to have a consistent non-directional distance between two detections D_i_ and D_j_, we can define it as

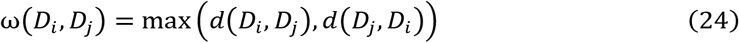

The distance function ω in Equation 24 can also discriminate the condition that a small detection is fully overlapped within a large detection. The directional distance from the small detection to the large one will be zeros, but if we measure it using the counter direction, it will be large.

#### Minimum-cost circulation framework

Current minimum-cost flow solvers are not efficient enough on large-scale datasets. These solvers utilize the strategy that pushes successive shortest paths until optimum (*27*) or binary searches the optimum given pre-determined flow amounts. For the scenario of cell tracking, the best time complexity is 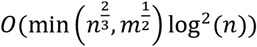 (*81*), where *n* and *m* are the numbers of cells and potential links, and the best real performance is to take several hours to track millions of cells. To solve the MAP problem mentioned above efficiently, we use CINDA (CIrculation Network-based Data Association) to solve the problem (*44*) (Fig. S5). This framework reforms the MAP problem as a minimum-cost circulation problem and maintains the same optimal solution. It has not only a superior 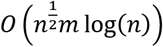 theoretical guarantee but also a tenfold practical efficiency improvement, which only takes minutes to track millions of cells and provides the feasibility of our iterative tracking strategy.

### Step 4: Error Correction

#### Cell under-segmentation and over-segmentation

Due to the ambiguity between adjacent cells or the intrinsic signal heterogeneity inside a cell, we may encounter segmentation errors, which mainly include two types. The cell under-segmentation indicates that there are indeed two cells (or more cells) contained in one detection. We wrongly merged the two cells. Cell over-segmentation is just the opposite, in which we wrongly split one cell into two (or more) detections. These two types of errors result in the same consequence, which is that two detections will have the same spatially closest neighbor in the adjacent frame. These errors have a clear difference from the normal cell division. For cell division, the two resultant cells will show clear spatial dis-connectivity, while for the segmentation errors, the two detections are tightly connected with each other. In our iterative framework, these segmentation errors are processed by module one.

We first designed a new network based on minimum-cost circulation, which links detections across time while allowing detections to split or merge. As is shown in Fig. 3D, a cell is mistakenly split as two detections in frame *t*+1, while the segmentation is correct in frame *t* and *t*+2. We add an arc from s to h_1_ the post-node of detection 1 and thus h_1_ can have at most two units of input and output flow. By the same token, we add one arc from o_4_, the pre-node of detection 4 to s, which allows at most two units of flow going through it. Now detection 2 and 3 can both be linked to detection 1 and also can both link to detection 4. The over-merged cells can also be processed with this design where the arcs are appended to the detection consists of two cells.

It is worthy to note that, we usually cannot determine a two-to-one linking is cell over-segmentation or under-segmentation directly from these two or three adjacent frames as evidence is still limited. Otherwise, we can directly force the over-split detections to be merged or an over-merged detection to be split. However, utilizing our new network design, the over-split or over-merged detections will be involved in a long trace covering dozens of frames. With the rich spatial-temporal relationships inferred from this trace, we can resolve the errors by re-merging and re-splitting the ambiguous detections.

We designed four criteria to determine the one-to-two and two-to-one linking is over-segmentation or under-segmentation. In practice, the four criteria will be applied sequentially as the confidence of the conclusions drawn from these them are different.

##### Voting from adjacent frames

The first criteria is to utilize the information from adjacent frames. For example, for the detection 1274 in the trace in Fig. S7A, the former three frames and the latter three frames all contain two detections. If we view the condition in one frame as a Bernoulli trial with over-segmentation probability 0.5. The probability of 6 out of 7 frames are over-split is less than 5%. Thus, we conclude that detection 1274 should be forced to split (Fig. S7B). Similarly, we can tell detection 3696 and 3697 should be forced to merge.

##### Voting from disconnected detections

The second criterion is based on the assumption that if a cell is over-split at some time point, the resultant two detections should be close and touched. Thus disconnected detections can be viewed as a piece of evidence indicating that there are two cells intertwined in this track. For example, for the track in Fig. S7A, the detection 55 and 78 are in the same frame but totally disconnected (Fig. S7B). Based on our assumption, we can conclude that the two-to-one linking temporally closest to the disconnected detections should be forced to split, which means the detection 1274 should be split. Note that, this splitting decision for detection 1274 has been made using our first criterion. In practice, the second criterion will only be applied if the first criterion failed to give a solid conclusion.

##### Voting from detections with similar size

We will conduct the third criterion if the former two both fail. The third criterion is based on the assumption that the territory size of a cell should be relatively consistent. For example, from t = 19 to t = 40, there is only one cell involved in the track with a relatively similar territory size. Among these 22 frames, we have two detections in 5 frames, while one detection in all other frames. With the same idea from our first criterion, the probability of 17 out of 22 frames are over-merged is less than 5%. Thus, we conclude that detections in the 5 frames are all over-split and should be forced to merge.

##### Try splitting

If all the previous criteria fail, we will try splitting the detection based on the gaps detected by principal curvature. If the resultant two detections have a high affinity score with the two detections in the former or latter frame, we will keep them. For example, for the detection 1742 in Fig. S7B (t=15), if we split it, the resultant two detections will have high morphological similarity with detection 1622 and 1623. Thus, we will choose to force detection 1742 to split. The reason we did this is to retrieve the false negative gaps when we conduct cell detection and segmentation. The gaps used here must be insignificant and missed in gap testing, otherwise we should have already split the corresponding detection.

With these four criteria, majority of the under-segmentation and over-segmentation cells should be corrected, the rest of which will be left for the next iteration when more evidence is accumulated. With such an iterative framework, we indeed are not afraid of making wrong decisions on the splitting and merging. As long as majority decisions are correct, the wrong decisions conducted will highly likely be corrected in the next iterations.

#### Cell missing

Fig. 3B shows a typical example of cell missing. Here we call the detection in the former frame as a parent of the missed detection and the one in the latter frame as a kid of the missed detection. The cell in frame t is missed while its parent and kid are both detected. Thankfully, our minimum-cost circulation-based data association framework has the ability to deal with this kind of cell missing.

Fig. 3C illustrates how it works. As is shown in Fig. 3B, there are two cells in the field of view indicated by different colors. The blue cell forms a track that consists of detection 2, 3 and 5. The orange cell has only two detections since its footprint in frame *t*+1 was missed. To continue the tracking of the orange cell, our network shown in Fig. 3C allows the linkages between detections beyond adjacent frames. Thus, though detection 1 has no appropriate candidate to link in frame *t*, it can jump over this frame and directly link to detection 4. In practice, we can adjust the number of frames that are allowed for jumping by a rough estimation of the missing rate.

Based on detection 1 and 4, we could have a reliable estimation of where the cell should be at time *t*+1 and retrieve it. The procedure is similar to our seed-based cell territory refinement. In practice, the seed region is set as the intersection between the parent detection and kid detection at the frame that missing happens.

It is worthy to note that the two modules in our framework focus on different aspects to refine the cell detection and segmentation results. To avoid making the system cumbersome, we did not allow jump when dealing with under-segmentation and over-segmentation problems in module one. Similarly, we did not allow detections to split or merge in module two when dealing with cell missing.

### Step 5: Post-processing

#### Tracklets association

In the iterative tracking and error correction stages, we control a reasonable FDR when designing the costs in Equation 20, while untracked outliers always exist and can lead to broken tracklets. To completely reconstruct the lineage, we execute the track stage one more time but consider the head and tail of a track only. Because most of the reliable motions have been successfully associated, the new distance distributions considering heads and tails only can have a larger variance, and hence the outliers with longer motions can be associated under the control of the same FDR. This step usually eliminates false negative associations without bringing more false positives since extra information from tracking results is utilized.

If a user would rather further complete the cell lineage than improve the tracking performance, another option is to associate cells compulsorily. In this way, tracklets are associated as far as possible under some loose conditions, such as the maximal motion distance. It can help some high-level biological researches that may not be so sensitive to tracking accuracy, such as the embryonic development at the tissue and organ level.

#### Division detection

Division detection is a even more tough task than cell tracking because of several reasons. First of all, it refers to one-to-multiple matchings, which is more difficult and cannot be included in most tracking methods. Secondly, the cell morphology has a great transition after division, which leads to a lower affinity score or matching probability. And the most important reason is that division detection has a extremely high accuracy requirement. For example, cells in one embryonic data has a 1% probability to divide and the accuracy of a division detection algorithm is 99%. In this case, the final reconstruction result cannot benefit from introducing the algorithm at all since it will bring more errors (1%) than corrections (0.99%).

Therefore, in our pipeline, divisions are not processed together with normal cell tracking but in the post-processing module (Fig. 1F). The philosophy behind is to focus on the key problem at once, where the association of the majority cell transitions can be processed first and filtered, and then a much smaller number of detection candidates can tolerate more errors but maintain the same performance. Besides, another advantage of this arrangement is that division detection can learn more information from the tracking results as well. For example, if the tail of one track is adjacent to two heads of other tracks in the next time point, it is likely to be a division. However, if all the three tracks are long and stable, the probability of being a division is larger than three short and unstable tracks, because the reason causing broken can be multifarious for the latter, such as bad data quality. Benefiting from the robust tracking algorithm and high-accurate results, it is possible to detect most divisions only using some simple rule-based decision tree.

There are three major rules to distinguish division and normal cells.

##### Rule 1

Every cell nucleus is a dense ball, but the nuclei group is sparse. We assume the fluorescent imaging of a cell nucleus is dense without too much texture or gaps inside, and is not touched with its neighbors since nuclei are usually at the center of a cell. Hence, if two nuclei are touched, manifested as one segmented instance with two isolated and similar components, they are very likely to be cells in division. However, since the correctness of the assumption varies between data, where some imaging data has stripe artifacts and cause gaps in nuclei, the rule is optional and may not work on all data.

##### Rule 2

If a stable track starts with a small-volume cell as the head, it is very likely to be a cell in division. A stable track means the track length is long enough, which implies the guarantee of the data quality and small probability to meet a broken. If the size of the head cell of the track is half of that of its candidate parent, the probability of division is also very high. Since a division always brings a new track, this rule works for most divisions if segmentation and tracking are good enough (Fig. S9B).

##### Rule 3

The migration of cells in the field of view follows Brownian motion with drift (Fig. S9C). During cell division, two new cells usually carry momentum of the same size but opposite direction, manifested as rapid movement of the two cells in opposite directions. If candidate cells move in the same direction, it is almost impossible for true cell division to occur. We use this information and assume that the coordinates of the dividing cells in a certain frame are *B*_*0*_, and the coordinates of the two candidate offspring cells are *B*_*11*_, *B*_*12*_, respectively, then we have:

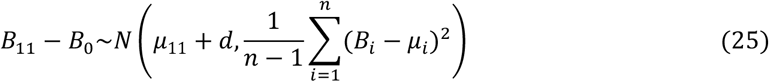

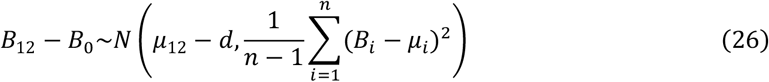

Among them, *d* is an additional drift caused by cell division. Due to the uncertainty of cell center markers and the influence of cell size during cell segmentation, the center position of cells follows the following distribution:

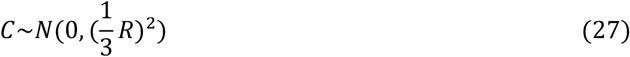

Taking into account the impact of the above two factors, there are

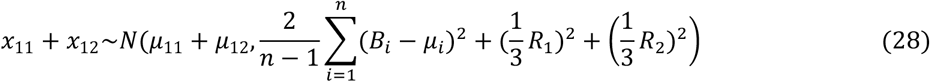

Note that *x* ∈ *R*^3^, assuming that each dimension follows the above distribution, then:

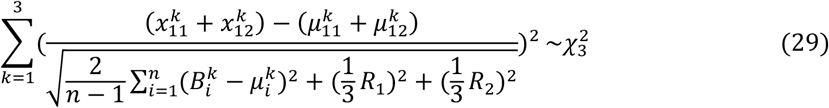

Based on the three rules, we design a decision tree (Fig. S9A). For every cell, first of all, we will check it has two isolated components or not, and then check if it is the head or second head of a track and satisfy several requirements simultaneously, such as track length and size ratio. Then we will try the best to assign the most matched parent or children if it passes all examinations.

#### Multi-angle stitching

To capture the whole embryo activity, some types of microscopes, such as light-sheet microscopes, records the specimen from different angles simultaneously, where cells may move out of the field of one view. There are two techniques to fully reconstruct the lineage: fusing the images or stitching the tracks. We adopted the latter for several reasons. Firstly, the image resolution is anisotropic, where the axial resolution is usually 3 to 10 times worse than the lateral resolution. Fusing images from different angles requires upsampling, means a huge increase of data size, and brings more pressure on memory and storage, but there is no information gain. Secondly, if one cell is recorded in multiple images, the data qualities are usually different, where the reason of multi-angle imaging is to guarantee good data quality in at least one image. Image fusion will average the data qualities, which is always worse than tracking the best one.

Oppositely, stitching tracks can not only consume nearly negligible computation and storage resources compare to image fusion but also make the best of all angles. We can get the approximate location correspondence after the spatial registration between all angles utilizing the approach discussed before. If a cell is highly-overlapped with another cell in a different angle, they are regarded as the same one. Then if two tracks have consecutively highly-overlapped cells, they are regarded as the same tracked and merged. Non-overlapped cells are preserved as much as possible, while long enough but not overlapped tails are also considered as divisions.

## Figures S1 to S15

**Fig. S1.**
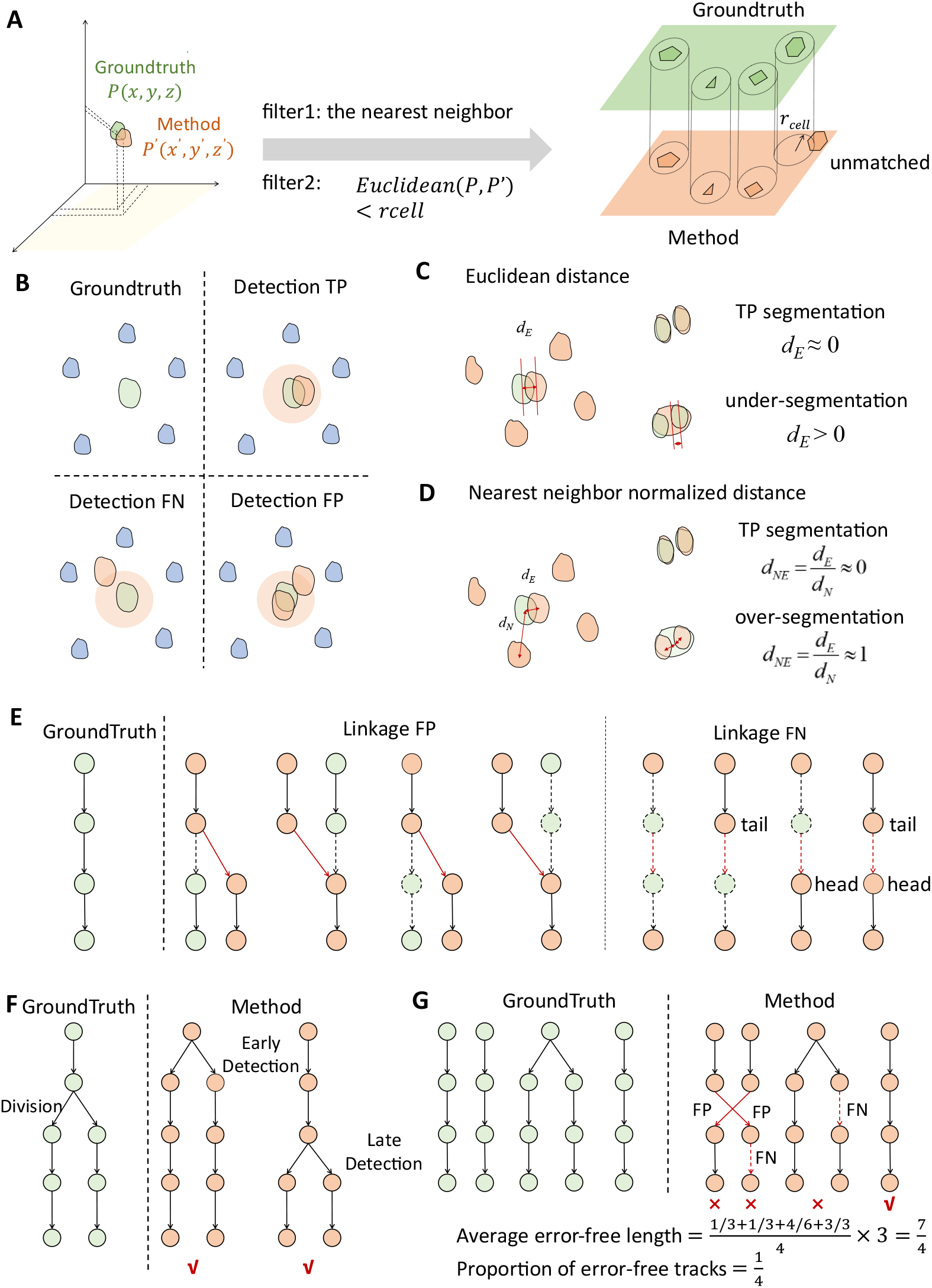
Evaluation metrics of cell tracking. (A) Cell matching between ground truth and method results. A cell detection is considered correct when a cell in the ground truth and a cell in the method results have an Euclidean distance that is less than the cell radius and are the nearest neighbors of each other. (B) Illustration of TP, FP, and FN cases in cell detection. A detection is considered TP when there is exactly one detected cell within a certain distance range of a ground truth cell. If there are more cells in this range, the excess cells are classified as FP errors. If there is less than one detected cell (i.e., none), it is classified as a FN error. (C) Euclidean distance metric for cell detection. This metric refers to the average of the nearest distances from each cell in the ground truth to any cell in the method results. If cell detection is completely correct, this metric is close to 0; if significant under-segmentation occurs, this metric will be large. (D) Normalized Nearest Neighbor Distance metric for cell detection. This metric is the Euclidean distance metric normalized by the average distance to adjacent cells in the method results. If cell detection is completely correct, this metric is close to 0; if significant over-segmentation occurs, this metric will approach 1. (E) Illustration of FP and FN in cell tracking. For a link between two consecutive frames in the ground truth, if the cell at at least one end of the link is incorrectly connected to a different cell, it is classified as a FP error. If this link is missing, whether due to a missing cell detection or a track break, it is classified as a FN error. (F) Evaluation correction for division events. Since cell division is a long-duration process and annotation errors may occur, we adopt a lenient evaluation strategy for divisions that occur earlier or later than in the ground truth, meaning they are not counted as FP or FN errors. (G) Illustration of the calculation for average error-free length and proportion of error-free tracks. We define these two metrics to characterize the continuity of the tracks.

**Fig. S2.**
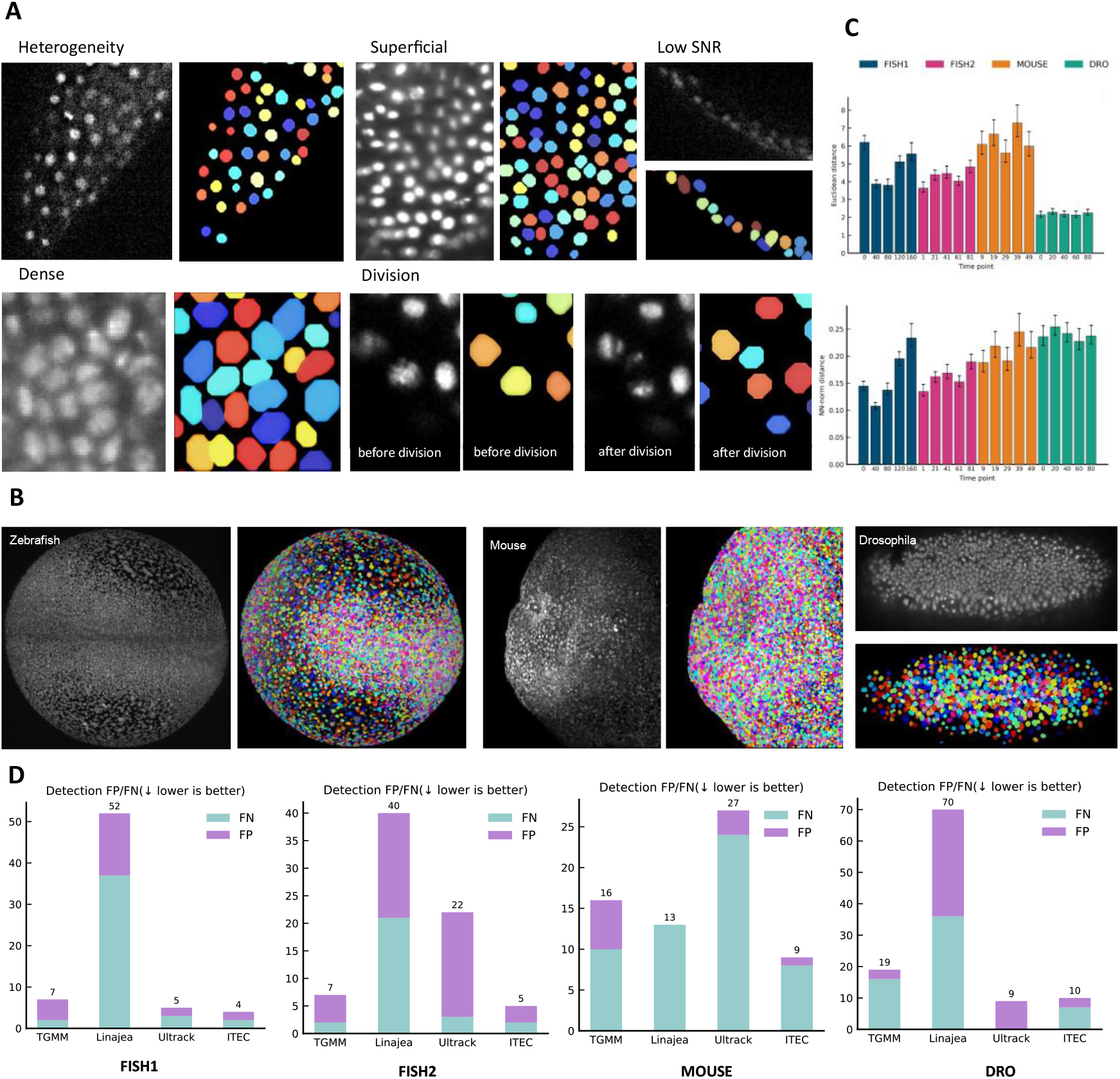
Robust cell detection in complex scenarios. (A) Local view of cell segmentation in different scenarios based on ITEC. ITEC achieves high-level segmentation under conditions of heterogeneity, superficial, low SNR, dense, and division. (B) Projection of cell segmentation for different species based on ITEC. The raw image uses the maximum z-axis projection, while the segmented image uses the nearest z-axis projection. (C) Average Euclidean distance (pixels) and normalized nearest-neighbor distance (error bars, 10^th^ and 90^th^ percentile confidence intervals). Both the Euclidean distance and the normalized nearest-neighbor distance are very low. (D) Detection FP/FN of ITEC and peer methods based on 500, 500, 250, and 500 annotated cells, respectively on the four datasets.

**Fig. S3.**
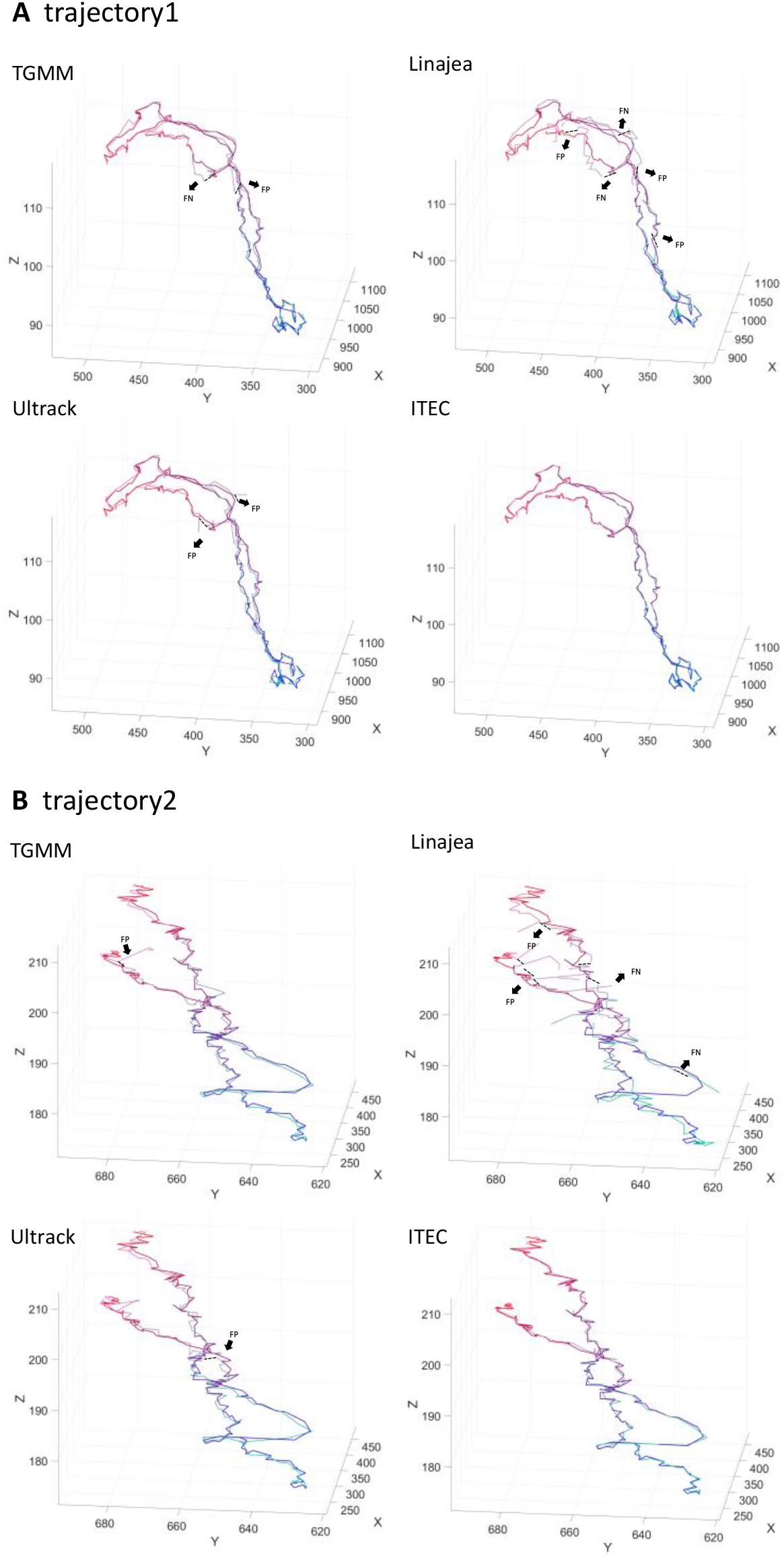
Comparision of tracking errors between ITEC and three peer methods. Track 1 and Track 2 are derived from the FISH1 and FISH2 datasets, respectively. Lines ranging from blue to red represent the ground truth, while lines ranging from cyan to magenta represent the algorithm results. Erroneous tracks are marked with dashed lines. Only the closest points of the algorithm’s coordinate points that match the ground truth are displayed.

**Fig. S4.**
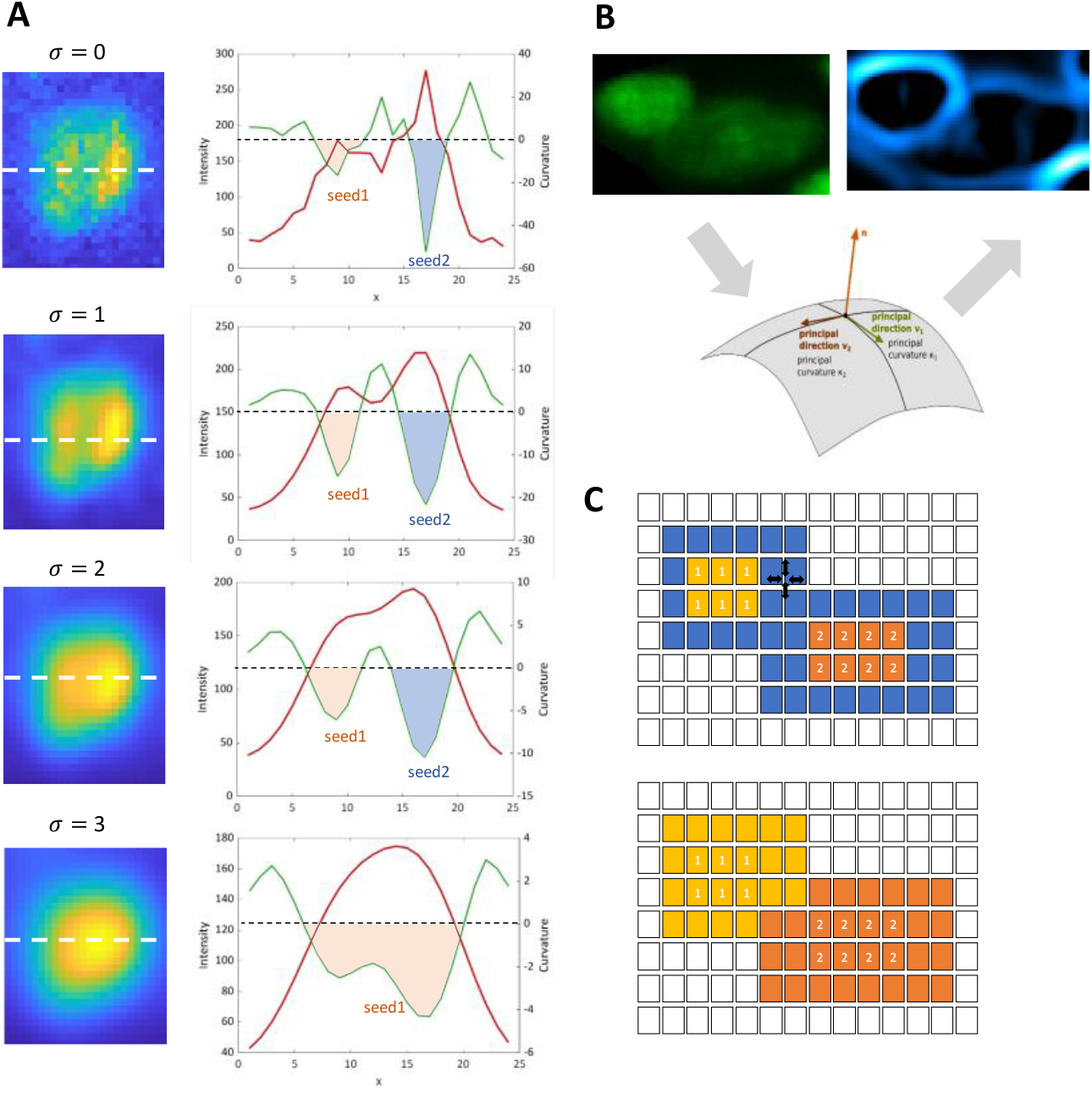
Some principles of ITEC cell segmentation (A) The high-order information of a typical cell, showing changes in cell brightness and curvature at different filtering scales. When the filtering scale is small, the cell is over-segmented into two cells. When the filtering scale is large, the over-segmentation is resolved, and the intensity of cell has only one peak. (B) Raw data and its curvature map. On the left is the original data, where two cells are squeezed together and it is difficult to determine their boundaries through simple grayscale information; On the middle is the geometric representation of principal curvature. On the right is the curvature map, and the boundary is clearly visible. (C) Seed regions. The yellow and orange colors on the left represent the seeds of two cells in (B), the blue color represents the area to be grown, and the right figure represents the well grown boundary.

**Fig. S5.**
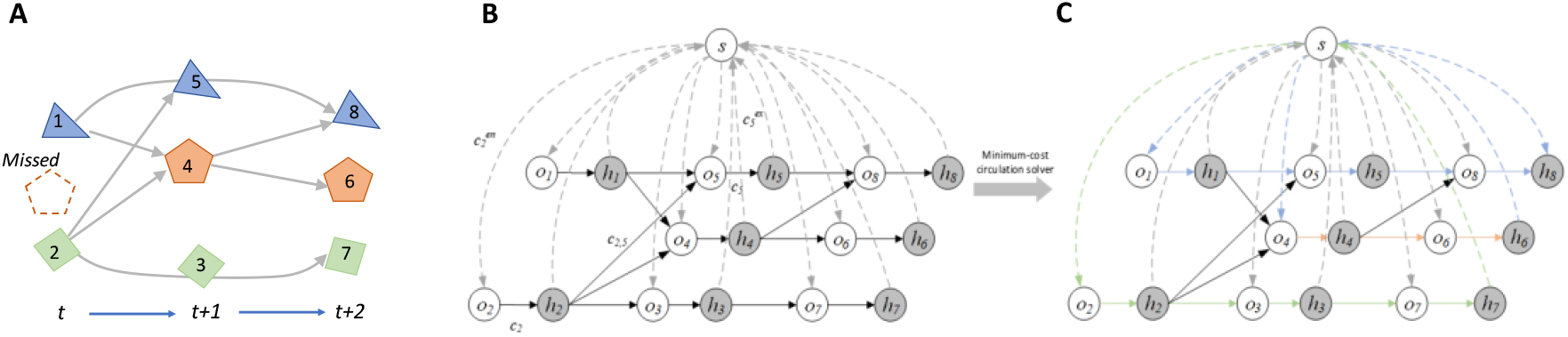
Minimum-cost circulation framework (A) Objects detected across three consecutive frames. The first frame includes two detections, with one missed detection indicated by orange dashed lines. The lines connecting detections represent possible associations, each assigned a certain cost. In total, three distinct tracks span these frames. For instance, detections 2, 3, and 7 belong to the same track and should be linked together. (B) The minimum-cost circulation formulation for MOT problem. Each detection *x*_*i*_ is represented by two nodes: a pre-node *o*_*i*_ and a post-node *h*_*i*_. A dummy node *s* is connected to all pre-nodes, and all post-nodes are linked back to *s*; these edges are shown as dashed lines. In this circulation network, flow conservation is maintained at every node, ensuring the dummy node’s excess flow remains zero. (C) The results from the proposed minimum-cost circulation framework. Three tracks are formed and they are shown with the same color as in (A).

**Fig. S6.**
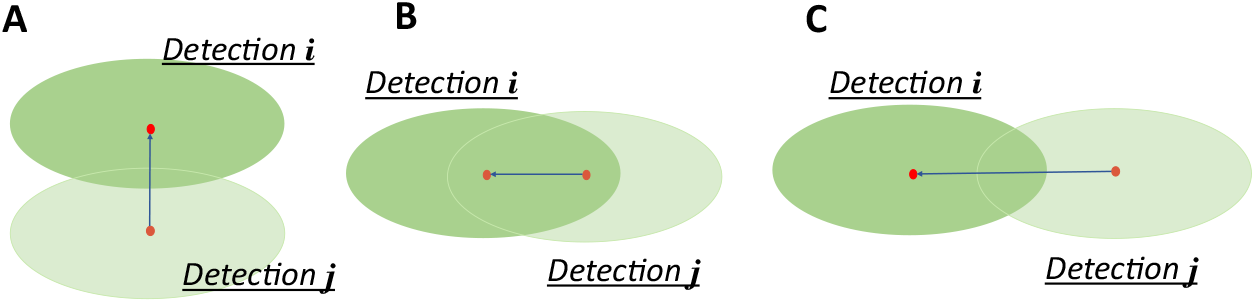
Illustration of the drawbacks of overlapping ratio and center distance. Here detection *i* and *j* are adjacent cell regions at different time points. If we want to measure their similarity, the desired score should be able to discriminate these three conditions as similarity (B) (A) (C). If we measure them using overlapping ratio, (A) and (C) become equal, while (A) and (B) will be equal if we use center distance.

**Fig. S7.**
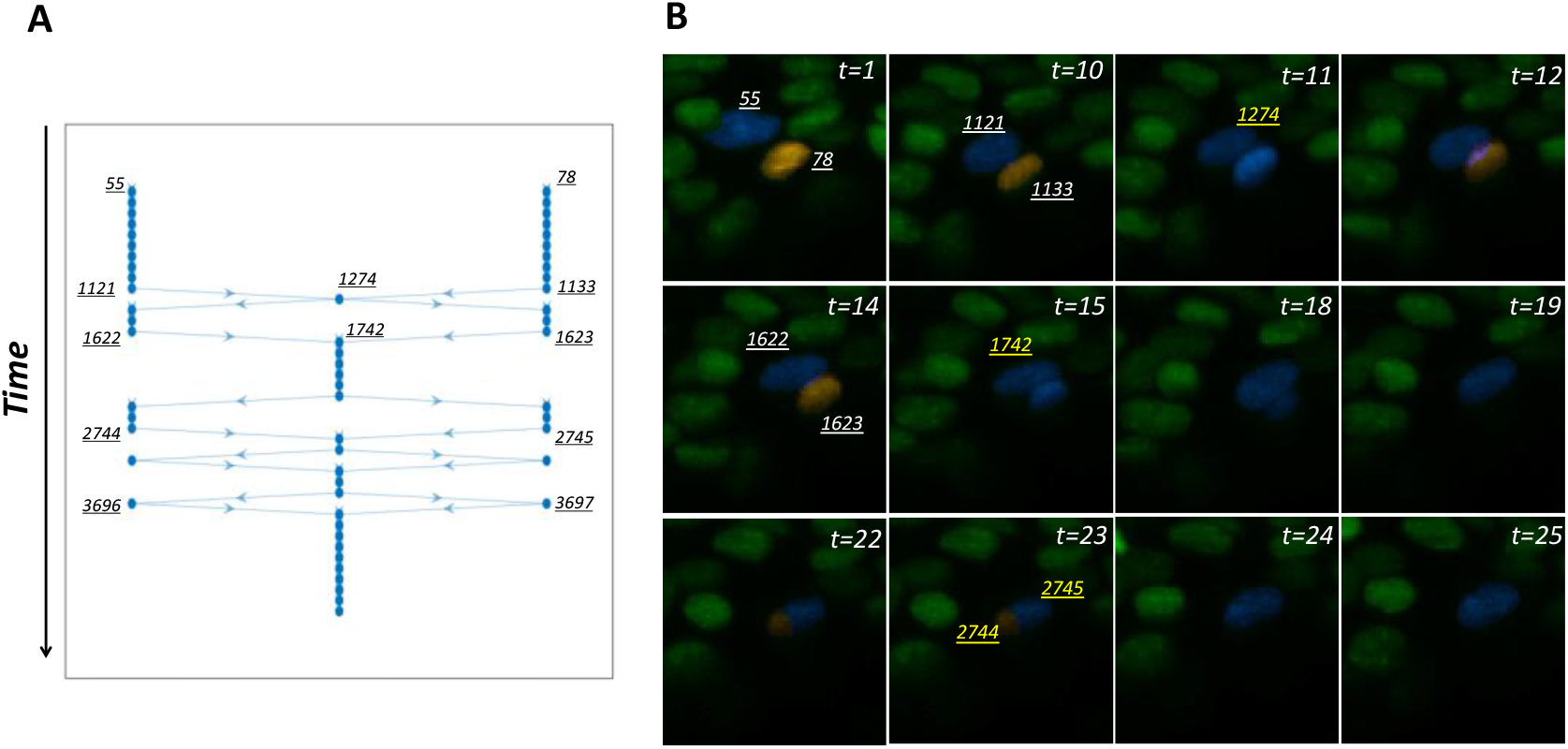
Example of over segmentation/under segmentation correction module (A) A typical error track. Each node represents a detection, and there have been several instances of over segmentation and under segmentation errors in the track. Our method will add one to two or two to one connections before and after nodes 1274, 1742, etc., and make subsequent judgments. (B) The original image corresponding to this example. The typical segmentation errors (yellow ids) and the corresponding segmentation results in adjacent frames. Totally two cells are involved in this track. One of them fades away at time point 18-19.

**Fig. S8.**
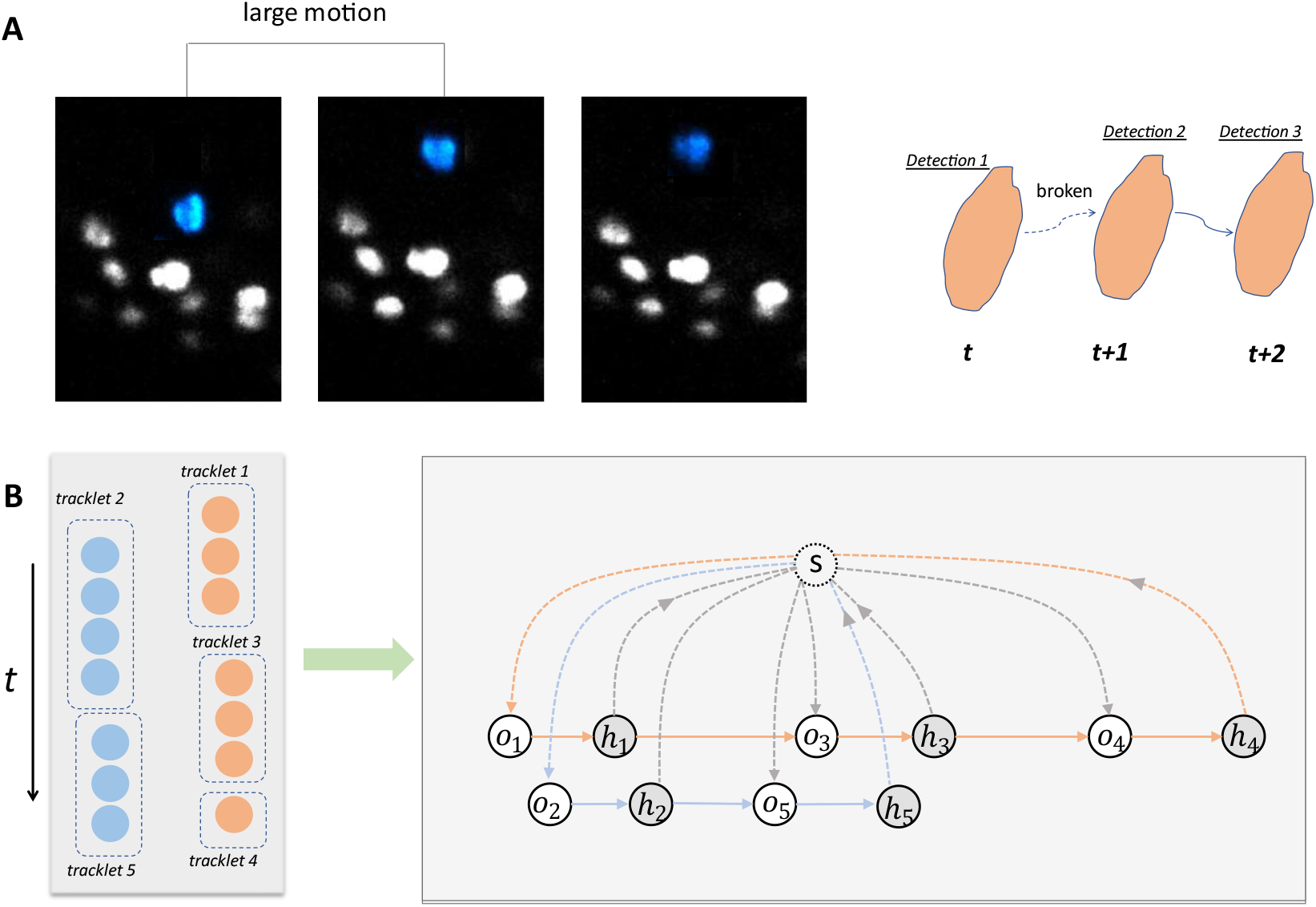
Tracklets association. (A) Typical track fracture. Due to the possibility of large-scale movement of cells, some associations may be disconnected during initial tracking. (B) Tracklets association network. We update the association cost based on the tracklets and use the minimum-cost circulation framework for new association.

**Fig. S9.**
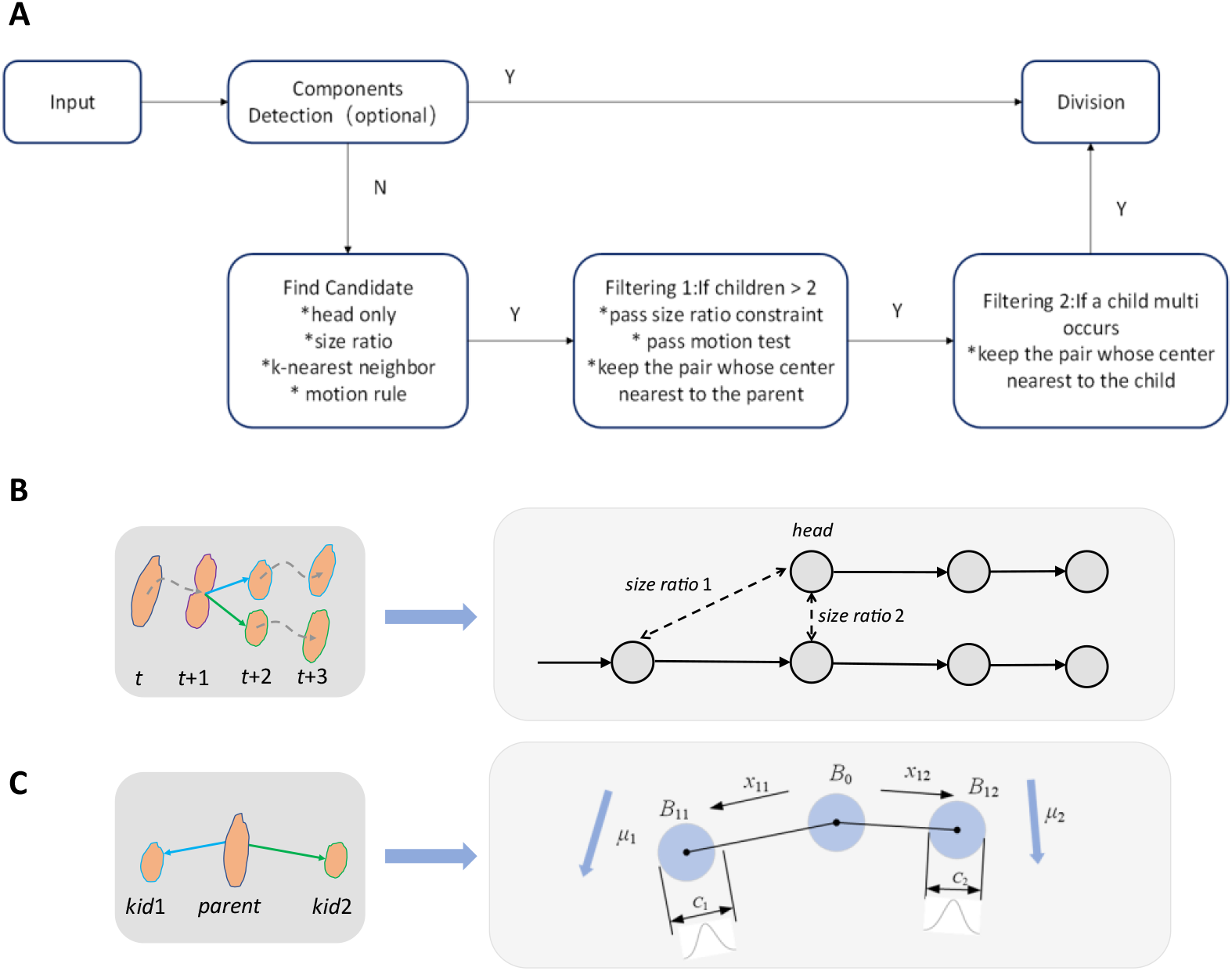
Principle of cell division module. (A) The pipeline of division detection. Firstly, for every cell segmentation result, it is reasonable but optional to treat a segmentation with two isolated components as divisions, under the assumption that cells are dense and integral. Then, for every candidate children, we will test it is a possible division or not. A candidate children must be the head of a track and satisfy the size ratio and motion rule requirements. It also need to be one of the nearest neighbor of its parents. For one parent, it is possible to have more than two candidate children. If so, we will check if there is a gap between any pair of two children, and only keep the pair whose center is nearest to the parent. Certainly, one child may also have multiple candidates as well. If so, we will follow the same strategy and only keep the nearest parent. The two steps can find most divisions. (B) Division candidate filtering. The child candidate needs to be the head of a fracture track and meet certain size constraints. (C) An example of cell division. *B*_0_ represents the parent cell, *B*_11_ and *B*_12_ represent the daughter cells. The two daughter cells have both the motion component caused by overall drift and the momentum generated by division.

**Fig. S10.**
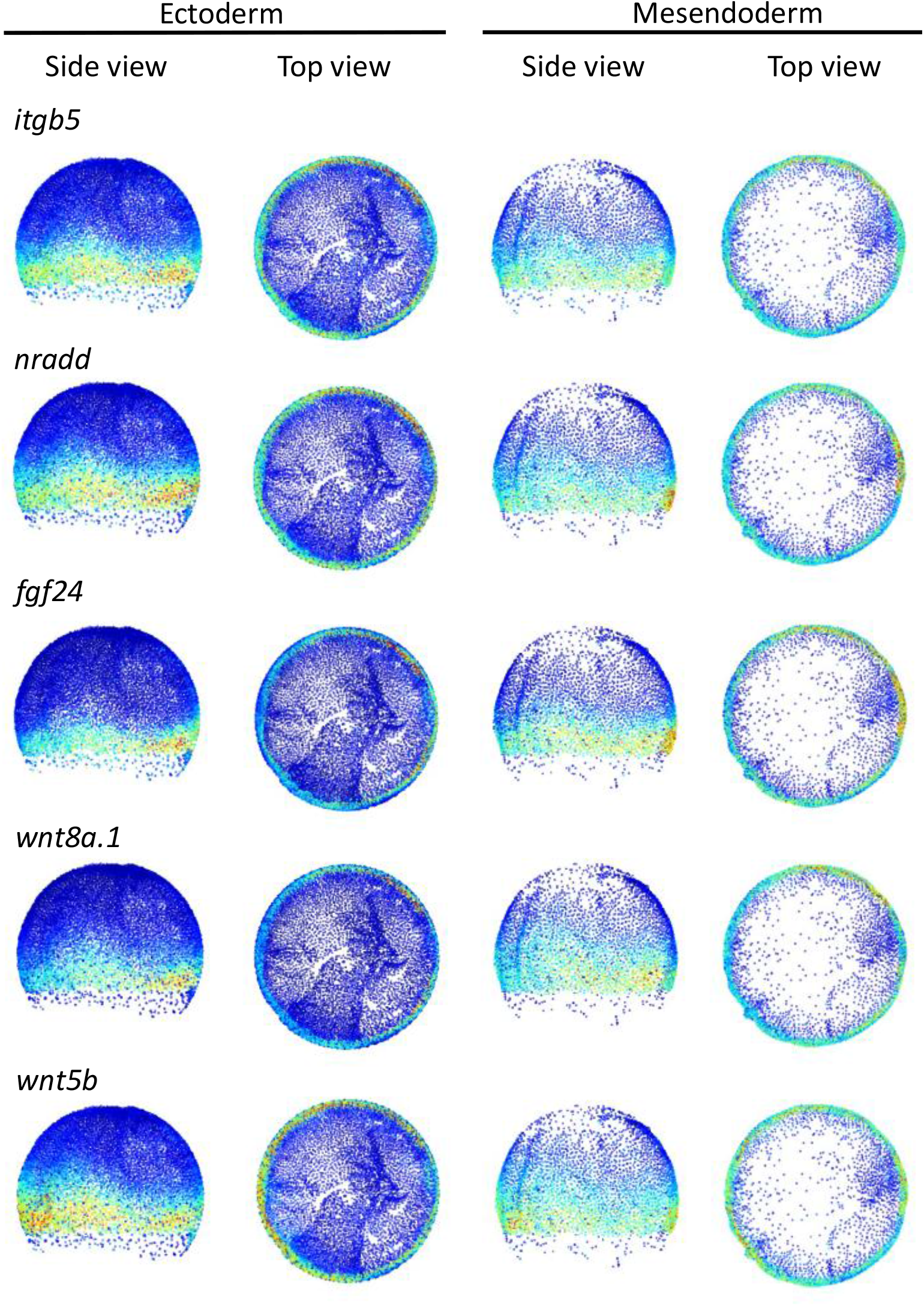
The expression of the 5 genes most related to cell migration velocity.

**Fig. S11.**
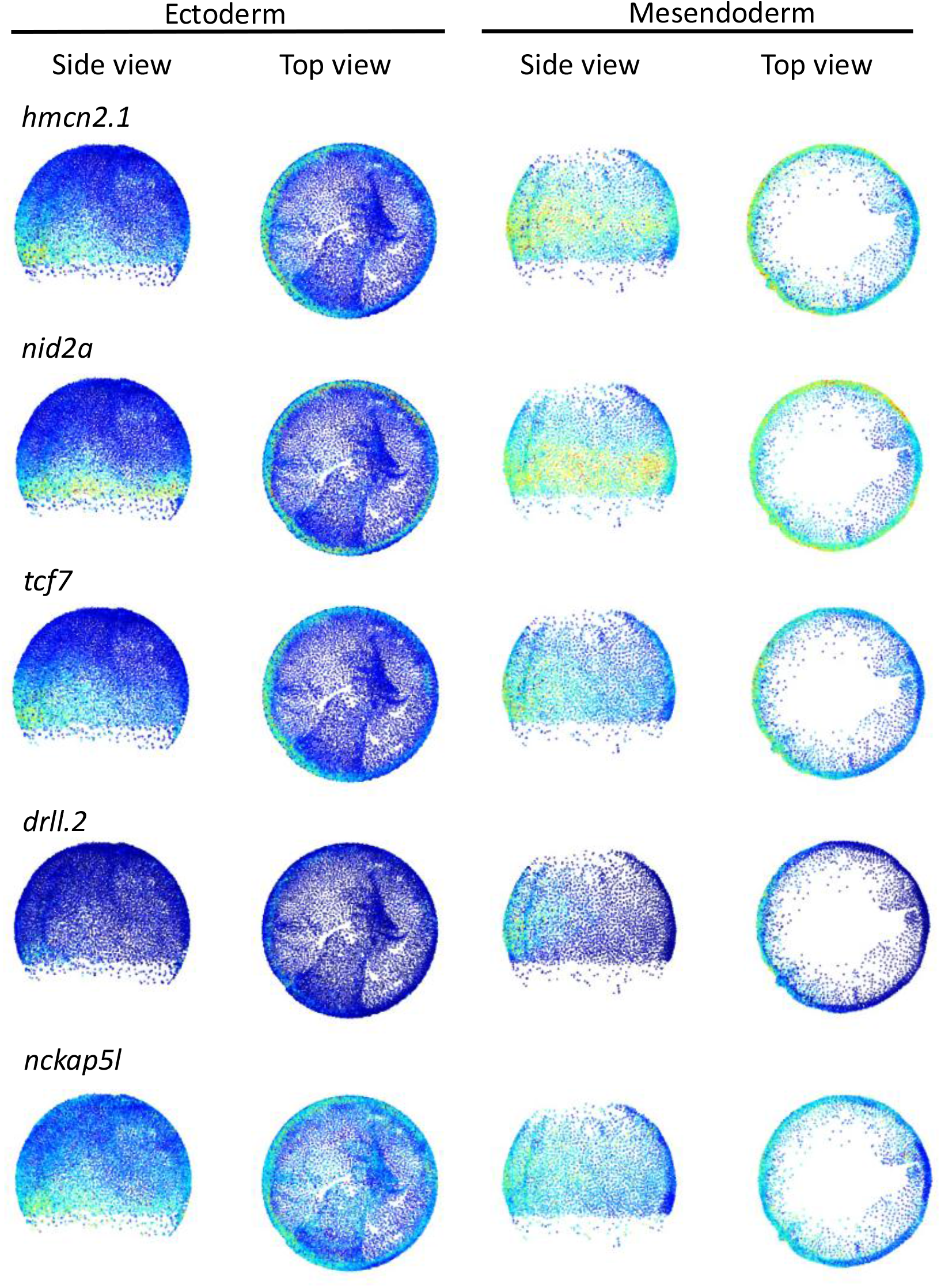
The expression of the 5 genes most related to cell motion variance.

**Fig. S12.**
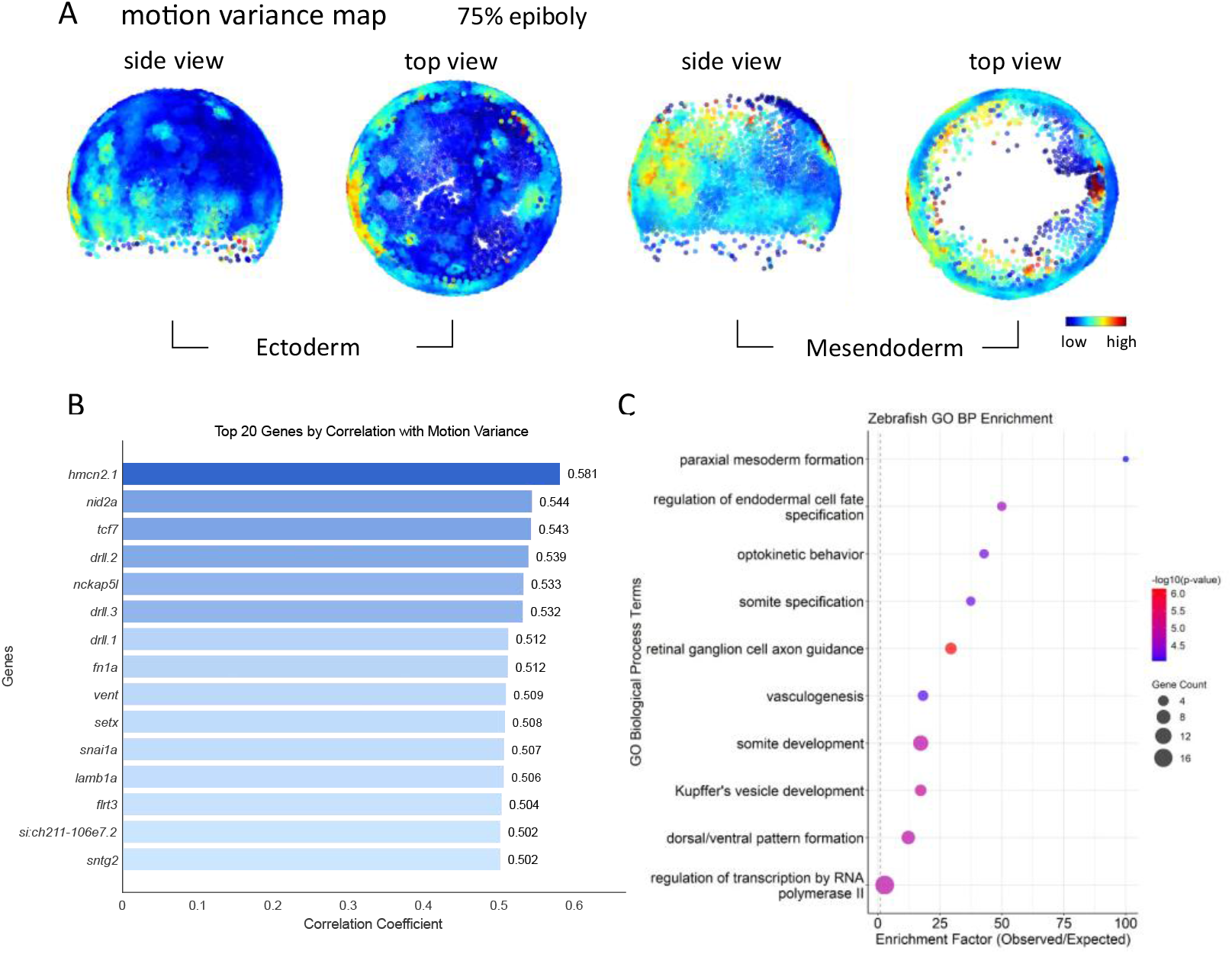
Analyze of cell motion variance at 75% epiboly. (A) Cell motion variance of ectoderm and mesoderm at 75% epiboly. (B) GO term enrichment of top 90 genes positively correlated with motion variance. (C) Top 20 genes most strongly positively correlated with motion variance and their correlation coefficients.

**Fig. S13.**
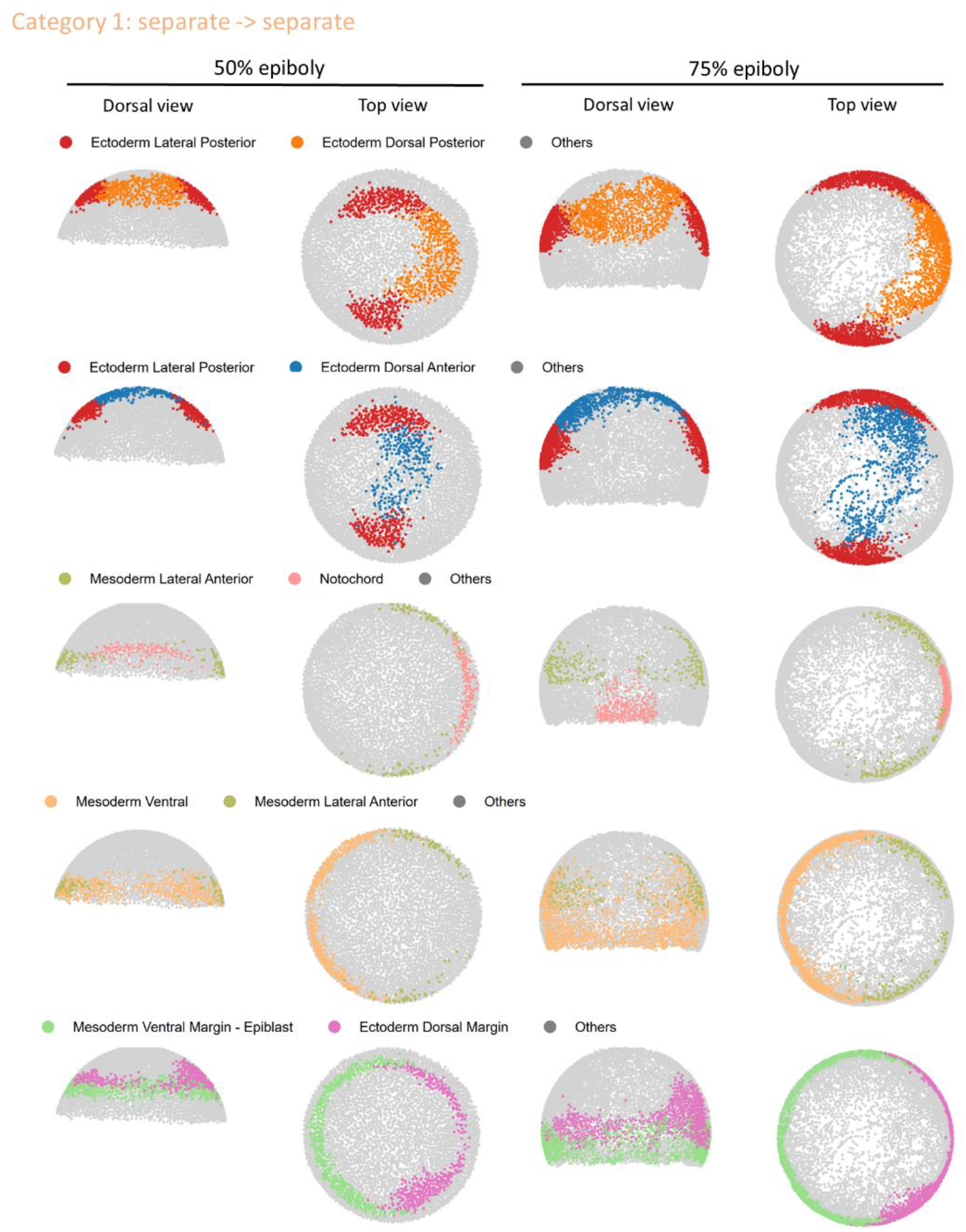
Examples of separate category boundaries maintained from 50% to 75% epiboly.

**Fig. S14.**
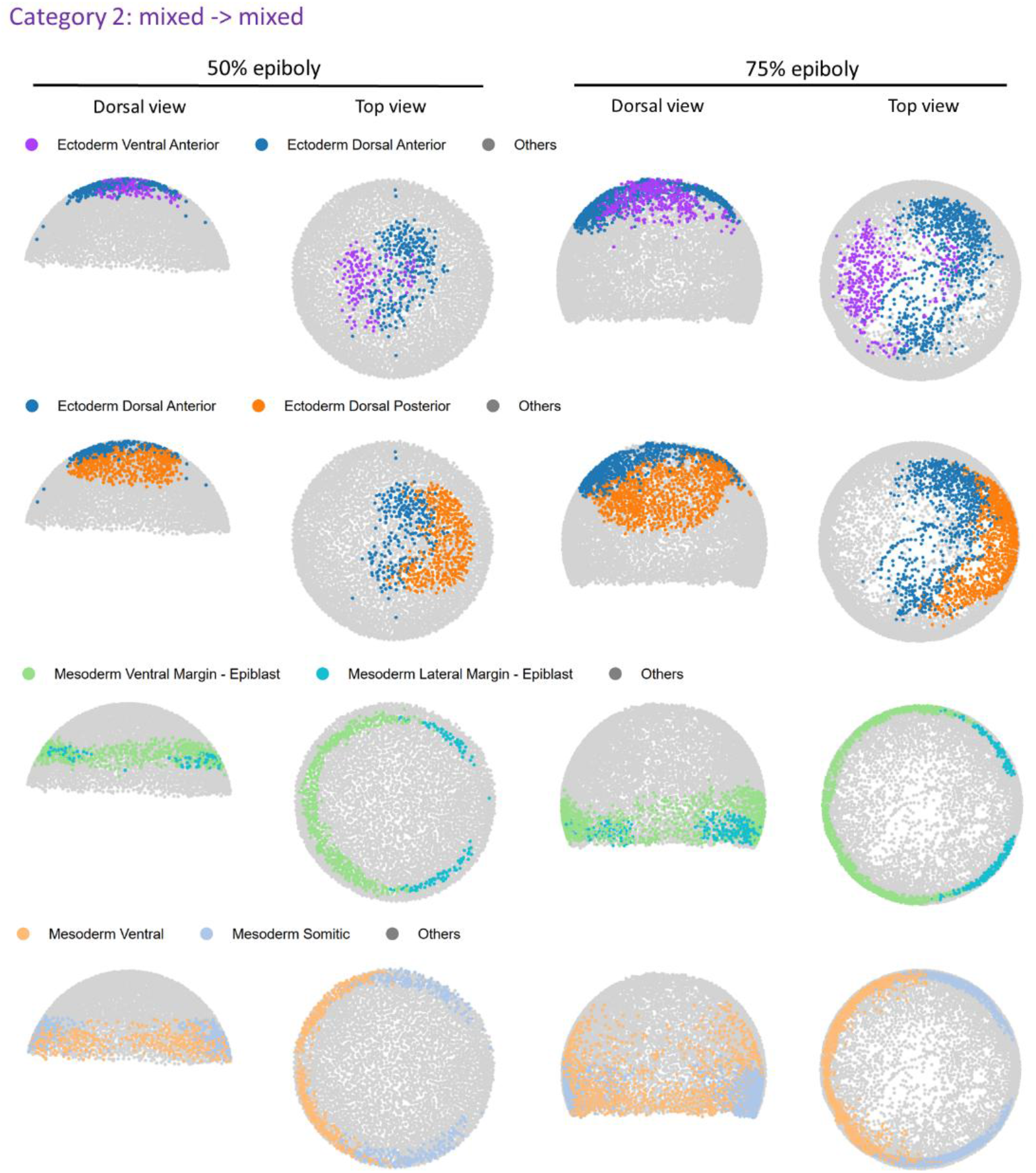
Examples of persistently mixed category boundaries from 50% to 75% epiboly.

**Fig. S15.**
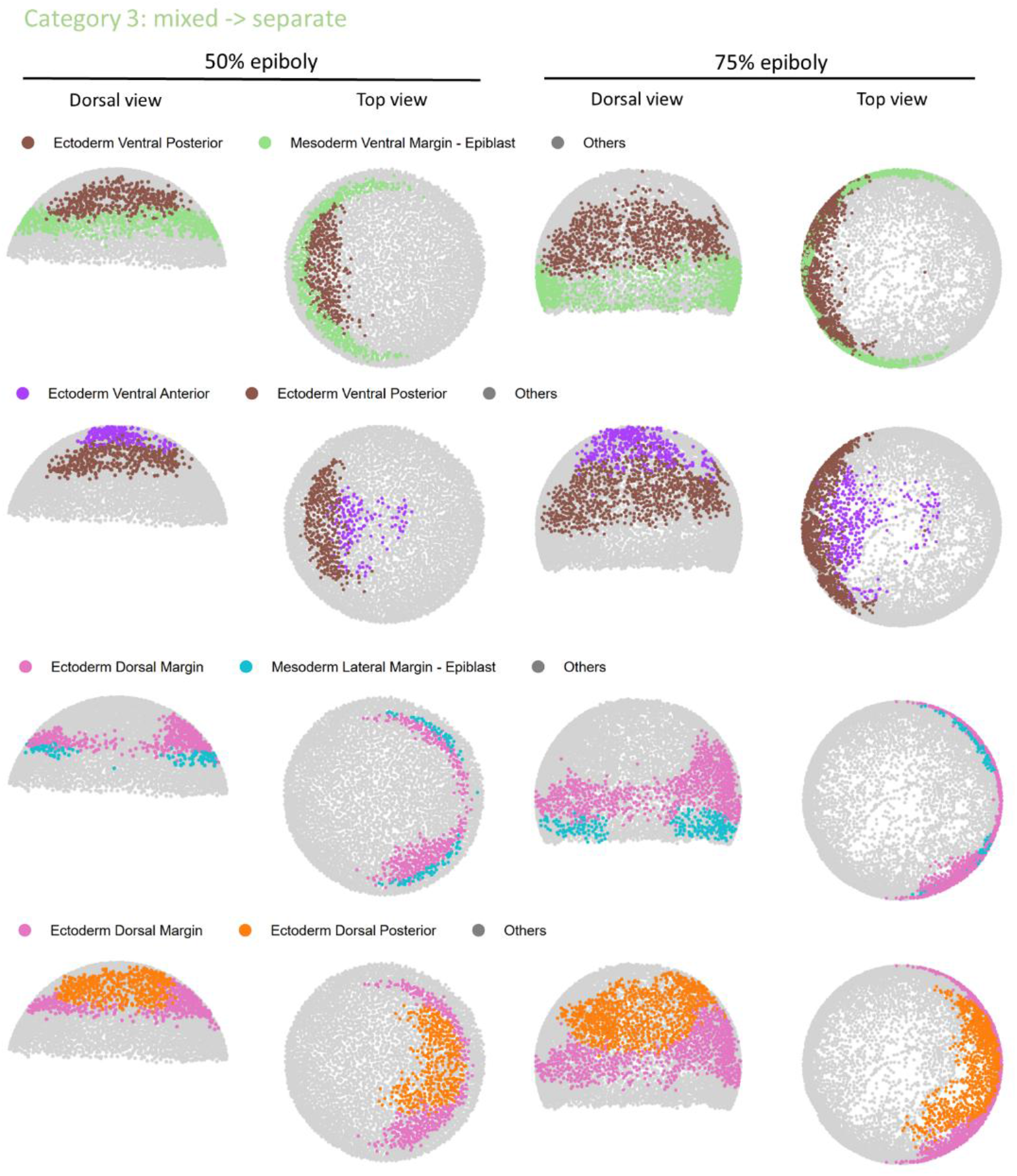
Examples of category boundaries transitioning from mixed to separate between 50% to 75% epiboly.

## Tables S1 to S3

**Table S1.** Peer method comparison.

| Methods | Training | Detection | Tracking |
| --- | --- | --- | --- |
| TGMM | Unsupervised | Watershed | Gaussian mixture model |
| Linajea | Supervised | U-Net | U-Net + ILP |
| Ultrack | Unsupervised | Multiple | UCM + ILP |

**Table S2.** Embryonic development datasets and annotations.

| Name | Animal | Image size | Time point | Cell (roughly) | Annotation |
| --- | --- | --- | --- | --- | --- |
| FISH1 | Zebrafish | $1920 \times 1920 \times 180$ | 192 | 1,590k | 32, 946 |
| FISH2 | Zebrafish | $1818 \times 1792 \times 253$ | 100 | 1,575k | 6, 841 |
| MOUSE | Mouse | $2169 \times 2048 \times 988$ | 50 | 815k | 2, 345 |
| DRO | *Drosophila* | $730 \times 320 \times 30$ | 100 | 66k | 2, 423 |

**Table S3.**
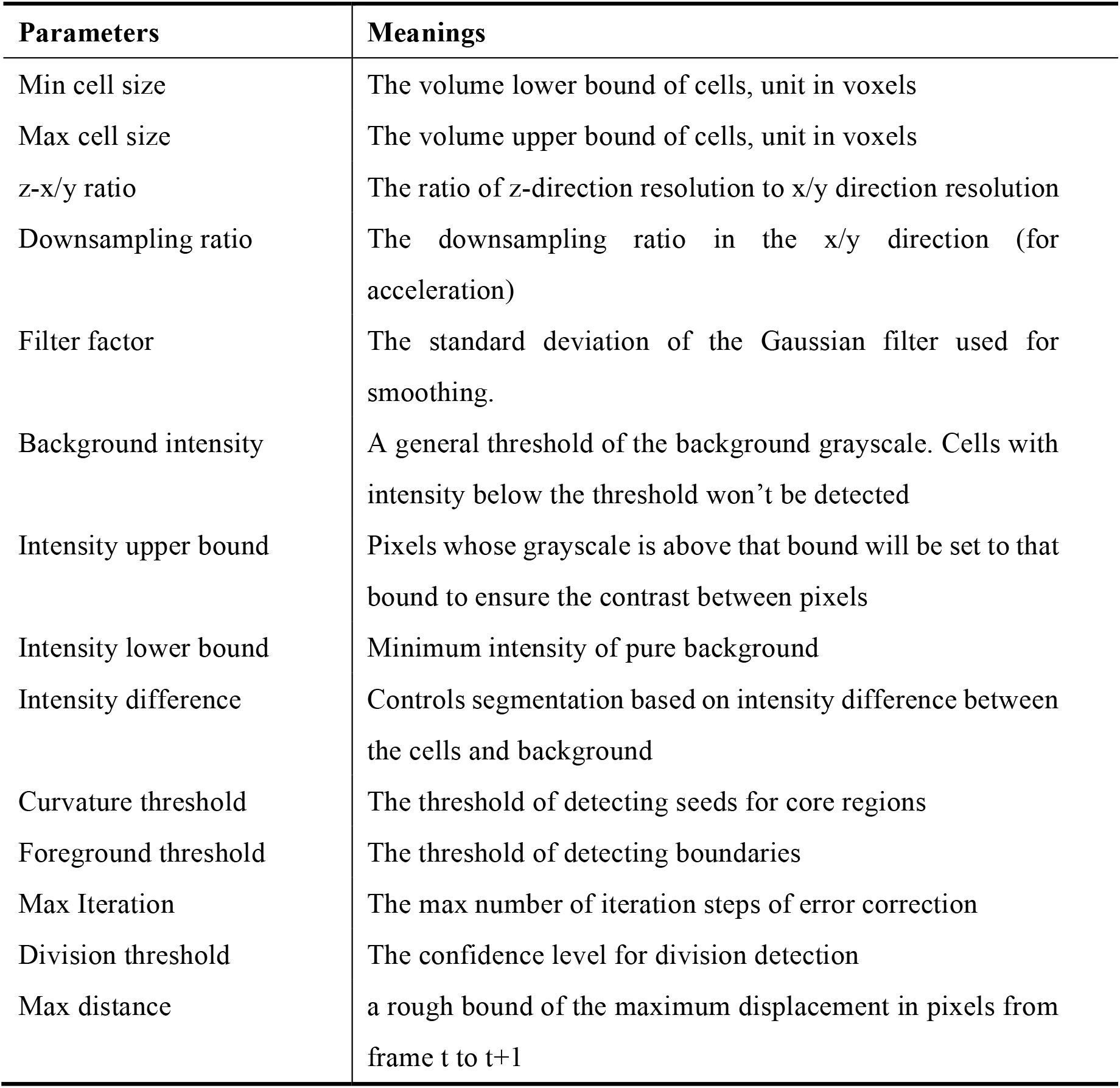
Some parameters in ITEC pipeline.

| Parameters | Meanings |
| --- | --- |
| Min cell size | The volume lower bound of cells, unit in voxels |
| Max cell size | The volume upper bound of cells, unit in voxels |
| z-x/y ratio | The ratio of z-direction resolution to x/y direction resolution |
| Downsampling ratio | The downsampling ratio in the x/y direction (for acceleration) |
| Filter factor | The standard deviation of the Gaussian filter used for smoothing. |
| Background intensity | A general threshold of the background grayscale. Cells with intensity below the threshold won't be detected |
| Intensity upper bound | Pixels whose grayscale is above that bound will be set to that bound to ensure the contrast between pixels |
| Intensity lower bound | Minimum intensity of pure background |
| Intensity difference | Controls segmentation based on intensity difference between the cells and background |
| Curvature threshold | The threshold of detecting seeds for core regions |
| Foreground threshold | The threshold of detecting boundaries |
| Max Iteration | The max number of iteration steps of error correction |
| Division threshold | The confidence level for division detection |
| Max distance | a rough bound of the maximum displacement in pixels from frame t to t+1 |

## Movies S1 to S5

Please see https://cloud.tsinghua.edu.cn/d/02fbca1ceceb490b814d/.

**Movie S1** An example of cell tracking using ITEC

**Movie S2** High-accuracy tracking using ITEC

**Movie S3** Long-term lineage reconstruction of zebrafish embryo

**Movie S4** Application of ITEC: Fate mapping

**Movie S5** Application of ITEC: Revealing the zebrafish somitogenesis

